# Increasing triacylglycerol formation and lipid storage by unsaturated lipids protects renal proximal tubules in diabetes

**DOI:** 10.1101/2021.09.07.459360

**Authors:** Albert Pérez-Martí, Suresh Ramakrishnan, Jiayi Li, Aurelien Dugourd, Martijn R. Molenaar, Luigi R. De La Motte, Kelli Grand, Anis Mansouri, Mélanie Parisot, Soeren S. Lienkamp, Julio Saez-Rodriguez, Matias Simons

## Abstract

In diabetic patients, dyslipidemia frequently contributes to organ damage such as diabetic kidney disease (DKD). DKD is associated with excessive renal deposition of triacylglycerol (TAG) in lipid droplets (LD). Yet, it is unclear whether LDs play a protective or damaging role and how this might be influenced by dietary patterns. By using a diabetes mouse model, we find here that high fat diet enriched in the unsaturated oleic acid (OA) caused more lipid storage in LDs in renal proximal tubular cells (PTC) but less tubular damage than a corresponding butter diet with the saturated palmitic acid (PA). Mechanistically, we identify endoplasmic reticulum (ER) stress as the main cause of PA-induced PTC injury. ER stress is caused by elevated cellular levels of saturated TAG precursors and to higher membrane order in the ER. The resulting cell death is preceded by a transcriptional rewiring of phospholipid metabolism. Simultaneous addition of OA rescues the cytotoxic effects by normalizing membrane order and by increasing the total TAG amount. The latter also stimulates the formation of LDs that in turn can release unsaturated lipids upon demand by lipolysis. Our study thus clarifies mechanisms underlying PA-induced cell stress in PTCs and emphasizes the importance of olive oil for the prevention of DKD.

## Introduction

Diabetic kidney disease (DKD; or diabetic nephropathy) is the most common complication of diabetes mellitus defined as diabetes with albuminuria or an impaired glomerular filtration rate (GFR) or both. It is the leading cause of end-stage renal disease, necessitating dialysis or transplantation. Despite improved management of diabetes, the number of DKD patients continues to rise causing an enormous health and economic burden worldwide. Classical histopathological features of DKD are glomerular changes such as podocyte hypertrophy and loss, glomerular basement membrane thickening, mesangial expansion and Kimmelstiel-Wilson nodules (Oshima *et al*, 2021). Often overlooked are the tubulointerstitial alterations, including peritubular fibrosis, that contribute to or even drive DKD progression (Bonventre, 2012). The recent success of sodium-glucose co-transporter-2 (SGLT2) inhibitors highlights the proximal tubule cell (PTC) as an important target in DKD therapy (DeFronzo *et al*, 2021).

Constituting more than half of renal mass (Park *et al*, 2018), PTCs reabsorb most of the solutes and proteins filtered by the glomerulus (Chevalier, 2016). While solute transport is carried out by dedicated transporters using ion gradients established by the Na^+^/K^+^-ATPase, protein uptake occurs via an endocytic machinery with low cargo specificity and unusually high capacity. The energy for both tasks is almost exclusively provided by mitochondrial β–oxidation of fatty acids (Bobulescu, 2010; Kang *et al*, 2015). The fatty acids are taken up from the blood side or by albumin that is partially filtered by the glomerulus and delivers bound fatty acids from the luminal side to the PTCs. Accordingly, in mice fed with a high fat diet or in individuals with type 2 diabetes, lipids accumulate predominantly in PTCs (Herman-Edelstein *et al*, 2014; Kang *et al*., 2015; Rampanelli *et al*, 2018), indicating that these cells may be equipped with a high capacity to take up and store lipids. Lipid overabundance, however, can lead to „lipotoxicity”, which is a main driver of kidney disease progression (Abbate *et al*, 2006; Bobulescu, 2010; Moorhead *et al*, 1982). This is particularly true in diabetic kidney disease (DKD), where albuminuria combined with dyslipidemia leads to a tubular overload of albumin-bound fatty acids (Bonventre, 2012; Zeni *et al*, 2017).

Within cells, lipid overabundance leads to enhanced triacylglycerol (TAG) synthesis and lipid droplet (LD) formation. This process occurs at the endoplasmic reticulum, where three fatty acids are consecutively added via esterification to the glycerol backbone beginning with *sn*-glycerol-3-phosphate. Once enough TAGs (and cholesterol esters) have been deposited between the ER lipid bilayer leaflets, LDs bud into the cytoplasm enwrapped by a phospholipid monolayer (Wilfling *et al*, 2014). In adipocytes, which are specialized in lipid storage, this is a physiological process. In other cell types, excessive LD formation is often a sign of impaired cellular homeostasis. A well-known example is hepatic steatosis that is featured by increased lipid storage in LDs of hepatocytes and often progresses towards liver fibrosis (Seebacher *et al*, 2020). Also in type 2 diabetes, lipid accumulation is a common feature in many organs contributing to insulin resistance. However, as free fatty acids (in particular, saturated ones) can activate pro-inflammatory pathways (Shi *et al*, 2006) or generate reactive oxygen species (ROS) upon excessive mitochondrial ß-oxidation, storing fatty acids in LDs could also prevent damage. Accordingly, it has been shown in neural stem cells that LDs can sequester polyunsaturated acyl chains, protecting them from the oxidative chain reactions that generate toxic peroxidated species and ferroptosis (Bailey *et al*, 2015; Dierge *et al*, 2021). Therefore, LDs can be damaging or protective depending on the tissue context.

In this study, we treated hyperglycemic mice with two different high fat diets, one enriched in butter (containing high amounts of palmitic acid (PA)) and one enriched in olive oil (containing high amounts of oleic acid (OA)). While the butter diet caused more renal fibrosis than the olive oil, less lipid accumulation was observed in the renal proximal tubules. By combining lipidomic, transcriptomic and functional studies, we find that PA induced rapid cytotoxicity by increasing the relative proportion of di-saturated TAG precursors in cellular membranes. This leads to ER stress which can be fully suppressed by co-incubating with OA. This protective effect is tightly connected with the formation of unsaturated phospholipids and the formation of LD that serve as a lipid reservoir to protect against lipid bilayer stress in the ER.

## Results

### A high fat diet enriched in saturated fatty acids causes tubular LD accumulation and kidney damage in diabetic mice

We wanted to study the effect of overloading PTCs with saturated and unsaturated fatty acids for DKD progression in mice. For this, we combined a low-dose streptozotocin (STZ) regimen, which destroys β-pancreatic islets and produces insulin deficiency, with two types of high fat diet (HFD) enriched in saturated fatty acids (SFA) and monounsaturated fatty acids (MUFA). Both HFDs contained 20 kcal% of protein, 35 kcal% of carbohydrates and 45 kcal% of fat, whereas the control diet contained 24 kcal% of protein, 58kcal% of carbohydrates and 18 kcal% of fat. The source of fat was butter for the SFA-HFD, olive oil for the MUFA-HFD and the standard soybean oil for the control diet (see also Table S1). The different diets were started at 7 weeks of age, and mice were followed for 20 weeks in four groups: while mice on control diet, SFA-HFD and MUFA-HFD received five consecutive daily injections of STZ (50 mg/kg) at 11 weeks of age, one group of mice on control diet was injected with the vehicle. Non-fasting hyperglycemia became apparent 4 weeks after STZ injection (Fig S1A). In addition, STZ-injected mice presented polyphagia and polydipsia as a sign of increased glycosuria (Fig S1B-C and Table S2). In addition, they showed increased levels of albumin in the urine (Table S2).

All three STZ-injected groups showed impaired weight gain over the entire experiment compared to vehicle-injected mice. Among the STZ-injected groups, no differences in body weight were observed except for the last two weeks when the mice fed a control diet decreased weight (Fig 1A). All STZ-injected mice had an extreme loss of epididymal white adipose tissue (eWAT), with some mice losing all their eWAT (Fig 1B). Concomitantly, plasma TAGs levels were increased in diabetic mice fed with MUFA-HFD and SFA-HFD (Fig 1C). Liver weight was only increased in mice fed with the two HFDs while all diabetic groups showed increased kidney weight (Fig 1B). We used periodic acid-Schiff (PAS) and Oil Red O (ORO) staining to determine whether the increase in kidney weight was due to ectopic fat deposition. Diabetic mice on both HFDs massively accumulated LDs in the kidney cortex. In the MUFA-HFD group, the percentage of kidney cortex stained with ORO was higher than in the SFA-HFD group. Individual LDs could be seen in tubules but not in glomeruli (Fig 1D-F and Fig S1D). Lipid accumulation was particularly evident in PTCs of the straight S2 or S3 segments but not of the convoluted S1 segment (Fig 1E).

**Figure 1.**
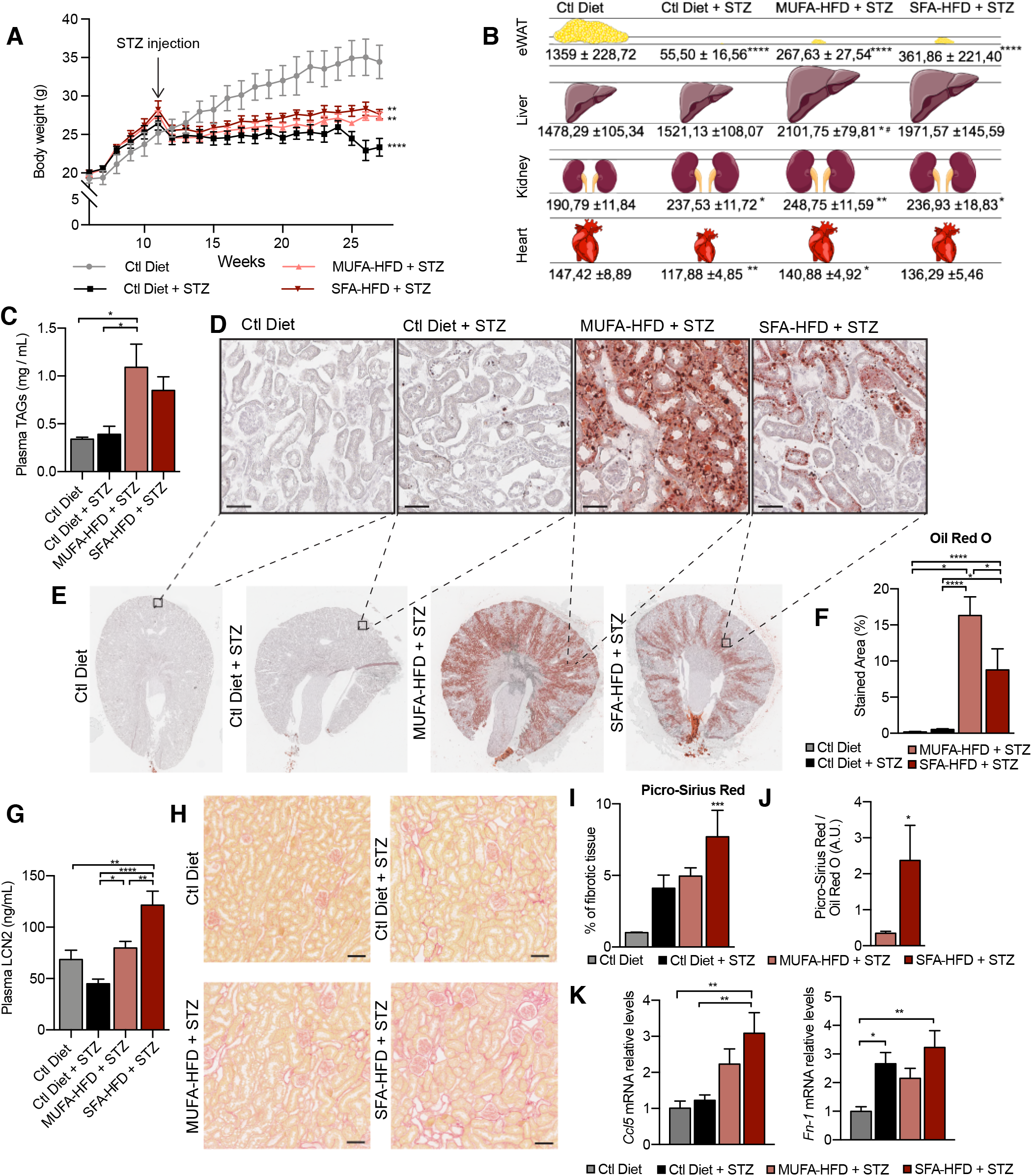
SFA-HFD induces more tubular damage than MUFA-HFD despite lower fat accumulation in mice. (A) Mice body weight throughout the experiment. The X-axis indicates the age of mice. (B) Schematic representation of tissue weight. (C) Plasmatic triacylglycerols (TAG) levels at week 16 after STZ injection. (D-F) Representative bright-field images of whole kidney sections (E) and cortex magnification (D) stained with Oil Red O. Quantification of the stained cortex area (F). Scale bars: 50*μm*. (G) Plasmatic LCN2 levels at week 12 after STZ injection. (H,I) Representative bright-field images of kidney cortex stained with Picro-Sirius Red (H) and the quantification of the fibrotic cortex area (I). Scale bars: 50*μm*. (J) Quantitative RT–PCR detection of *Ccl5* and *Fn-1* expression levels in mouse kidney cortex. (K) Fibrotic area detected by Picro-Sirius Red normalized by fat deposition measured by ORO in mouse kidney cortex. Data information: In (A,C,G,I-K), data are presented as mean ± SEM. *p<0.05, **p<0.01, ***p<0.001, ****p<0.0001; One-way ANOVA plus Holm-Sidak’s multiple comparisons test. In (B), data are presented as mean ± standard deviation. *p<0.05, **p<0.01, ****p<0.0001 vs Ctl; #p<0.05 vs Ctl Diet + STZ ; One-way ANOVA plus Holm-Sidak’s multiple comparisons test. (A,B,C,G,H,J,K,L), n=7 Ctl diet, n=8 Ctl Diet + STZ, n=8 MUFA-HFD + STZ, n=7 SFA-HFD + STZ.

To assess whether STZ and HFD treatments caused renal damage, we first studied circulating levels of the tubular damage marker LCN2. At 12 weeks after the STZ injection, LCN2 levels were higher in mice fed with SFA-HFD than in mice fed with MUFA-HFD (Fig 1H). To determine whether tubular damage resulted in fibrosis, we made use of Picro-Sirius Red staining to visualize collagen deposition. The staining revealed fibrotic areas in all diabetic mice groups but only in the mice fed SFA-HFD this increase reached statistical significance (Fig 1H-I). Moreover, when normalized to fat deposition, the increase in the fibrotic area became even more apparent in SFA-HFD kidneys compared to MUFA-HFD kidneys (Fig 1J). Increased mRNA levels of the profibrotic molecules *Ccl5* and *Fn1* in SFA-HFD kidneys supported the histological findings (Fig 1K). Finally, we performed immunohistochemistry to detect changes in the expression of the tubular damage marker KIM1 (Mori *et al*, 2021). We found strong apical KIM1 expression in renal proximal tubules in some of the STZ and SFA-HFD mice while only moderate in MUFA-HFD kidneys (Fig S1E).

Taken together, the STZ-induced diabetic mice recapitulated diabetic features, namely, hyperglycaemia, polyphagia, polydipsia, polyuria, glycosuria, albuminuria and lipodystrophy. While the MUFA-HFD diet favored the accumulation of LDs in the kidney, the overload with SFAs resulted in higher tubular damage and fibrosis.

### PA impairs cell viability in PTC culture models that can be rescued by OA

To better understand the molecular mechanisms driving SFA-mediated renal damage as well as MUFA-mediated renoprotection, we used induced renal epithelial cells (iRECs), which are proximal-tubule like cells directly programmed from fibroblasts (Kaminski *et al*, 2016). First, iRECs were treated with increasing concentrations of fatty acid-free bovine serum albumin (BSA) to determine the toxicity of albumin itself. However, none of the concentrations of BSA induced any cytotoxicity as measured by the uptake of Cytotox© dye using the Incucyte Live-Cell analysis system (Fig 2A). Next, we treated cells with BSA conjugated to PA, OA and combinations of both. PA treatment induced cell death after 8 hours in a concentration-dependent manner, while OA treatment did not affect cell viability at all. This was true for both sub-confluent and confluent cells. Importantly, co-treatment with OA completely rescued the cytotoxic effects of PA. Even when only 0.25mM OA was co-incubated with 0.5mM PA cell viability was fully restored (Fig 2B-D). To compare the effects with other PT cell culture models, we performed the same cytotoxicity assays on murine primary PT cultures as well as OK cells. In both cell models, similar effects on cell viability were observed when exposed to different doses of albumin-fatty acids (Fig 2E,F). Thus, it can be concluded that PA treatment causes cytotoxic effects in PTCs that can be rescued by OA.

**Figure 2.**
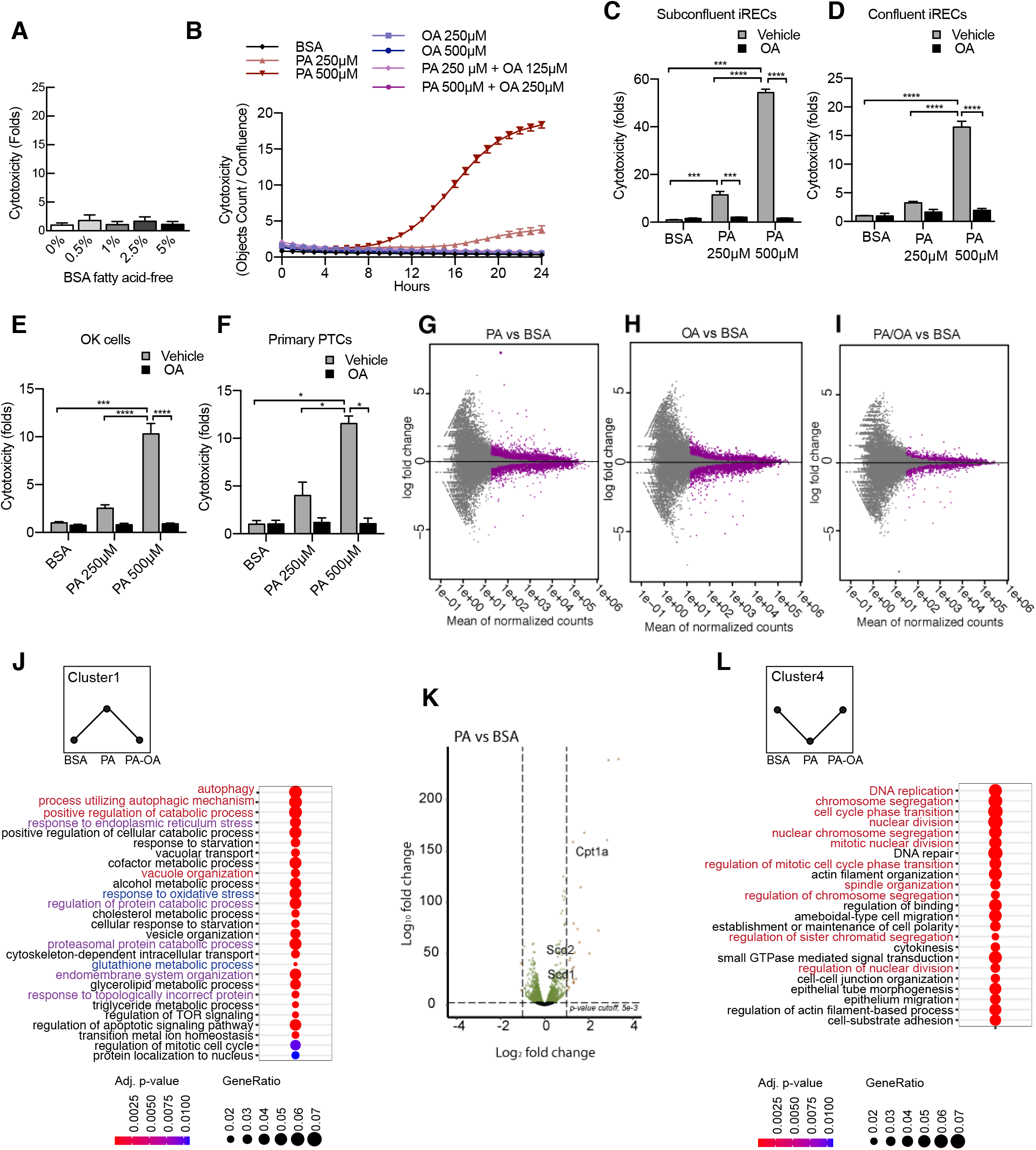
PA-induced cytotoxicity is blocked by OA in proximal tubular cells. (A) Cytotoxicity in iRECs at 24h treatment with increasing concentrations of BSA fatty acid-free. Fold representation of object counts / confluence. (B) Cytotoxicity throughout 24h in iRECs treated with several combinations of BSA-fatty acids. (C-F) Cytotoxicity in subconfluent (C) and confluent (D) iRECs, OK cells (E) and mouse primary proximal tubules (F) after 24h treatment with BSA, OA 250µM, PA 250µM, PA 250µM + OA 125µM, PA 500µM and PA 500µM + OA 250µM. Fold representation of objects counts / confluence. (G-I) MA-plots of differentially expressed genes from iRECs treated for 16h with BSA, PA 250µM, OA 250µM and PA 250µM + OA 125µM. Purple dots represent statistically significant changes. (J,L) GO for biological processes overrepresentation analysis using clusterProfiler. The 25 most significant terms were plotted for clusters 1 (J) and 4 (L). The size of the spheres corresponds to the number of genes included and the colour to the adjusted p value. (J) Biological processes related to autophagy coloured in red, related to oxidative stress in blue and related to ER stress in violet. (L) Biological processes related to cell proliferation coloured in red. (K) Volcano plot of PA vs BSA differential gene expression. Colored dots represent statistically significant changes. Data information: In (A-F), data are presented as mean ± SEM. *p<0.05, **p<0.01, ***p<0.001, ****p<0.0001; two-way ANOVA and Holm-Sidak’s multiple comparisons test. (G-L) significance was considered when adjusted p-value< 0.05 (DESeq2 based on negative binomial distribution). (A-E,G-L) n=3. (F) n=2.

### PA-induced cell injury elicits a unique transcriptional response

Next, we performed a comparative transcriptomic study on BSA-, BSA-PA-, BSA-OA- and BSA-PA/OA-treated iRECs using RNA sequencing. The differential expression analysis showed that the transcriptional response observed in BSA-PA and BSA-OA cells was much more pronounced than in BSA-PA/OA-treated cells (Fig 2G-I). As we were particularly interested in biological processes that were up- or downregulated by the addition of PA and reverted to normal levels when cells were co-incubated with OA, we clustered genes accordingly and applied gene ontology (GO) analysis. Cluster 1, that included genes upregulated by PA and normalized by OA co-treatment, proved to be strongly enriched in genes involved in oxidative stress, ER stress and autophagy (Fig 2J). Important lipid metabolism genes such as the mitochondrial fatty acid importer *Cpt1* and the fatty acid desaturases *Scd1* and *Scd2* also belonged to the top-regulated genes (Fig 2K). By contrast, cluster 4, featured by genes that were downregulated by PA and normalized by OA co-treatment, showed a clear enrichment of biological processes controlling cell proliferation (Fig 2L), confirming the observation that BSA-PA treatment slows down cell growth compared to the other three conditions (Fig S2A).

With the aim of identifying what transcription factors might be regulating these cellular responses, we mined our transcriptomic data using DoRothEA (Holland *et al*, 2020), a comprehensive resource containing a curated collection of transcription factors (TFs) and its transcriptional targets (see Materials and Methods). We obtained TFs whose predicted target genes associated with one or more of the five different conditions (PA vs. control, PA vs. PA/OA, PA/OA vs. control, OA vs. control, OA vs. PA/OA). Interestingly, several of the identified TFs whose activities were upregulated by PA and downregulated by OA (Fig S2B) have known roles in lipid metabolism (e.g. HNF4A, SREBF1) and ER stress (e.g. ATF4, ATF6). By contrast, several TFs whose activities were regulated in the opposite way function in cell cycle control (e.g. E2F4, MYC, FOXM1). Thus, the TF analysis confirmed the initial transcriptome data analysis by highlighting upregulated ER stress and lipid metabolism and downregulated cell cycle progression as the main response to PA.

### PA-induced ROS formation is blocked by OA cotreatment

To functionally validate the transcriptional responses, we focused on the cellular processes upregulated by PA (cluster 1). To study the oxidative stress response, we used dihydroethidium (DHE) to measure the generation of ROS. Superoxide levels were increased by PA treatment in a dose-dependent manner. When OA was added together with PA, the DHE staining returned to basal levels (Fig S3A-B). Next, we wondered whether the protective effect of OA on ROS formation was due to changes in mitochondrial activity. Using tetramethylrhodamine ethyl ester (TMRE), which is an indicator of mitochondrial membrane polarization, PA was found to increase mitochondrial activity while OA reduced it. The combination of PA with OA returned mitochondrial activity to basal levels (Fig S3C,D). As β-oxidation requires the import of fatty acids into the mitochondria by CPT1, which was upregulated by PA and OA, we used the CPT1 inhibitor etomoxir that was previously shown to decrease mitochondrial β-oxidation in iRECs (Marchesin *et al*, 2019). Etomoxir treatment increased the PA-induced cytotoxicity. However, OA was still protective in this condition (Fig S3E).

Together, these experiments showed that PA stimulates ROS generation and OA blocks it, validating the observed antioxidant response in the RNA-seq study. Since etomoxir boosted PA-induced cytotoxicity, increased mitochondrial fatty acid uptake and oxidation seems to be a beneficial response to PA overload. Yet, the protective effect of OA does not seem to involve these processes.

### Albumin-PA triggers an ER stress response that can be rescued by OA

Next, we examined the role of ER stress in response to the PA insult. ER stress can be activated by three different branches known as the IRE1, ATF6 and PERK branches (Walter & Ron, 2011). qPCR-based ER stress marker analysis of spliced *Xbp1 (sXbp1), Hspa5 (*also known as *Bip), Ddit3* (also known as *Chop*) mRNA showed that all three ER stress response pathways are activated by PA after 16h. Full suppression of ER stress activation was achieved by cotreatment with OA (Fig 3A). In order to study which of the ER stress branches contributed most to cytotoxicity, we co-treated BSA-PA iRECs with inhibitors of PERK (GSK2606414), IRE1a (4μ8C) and ATF6 (Ceapin-A7). Only the PERK inhibitor slightly reduced the cytotoxicity caused by PA, suggesting that the PERK-eif2a-ATF4 axis may contribute to the cell death process (Fig 3B).

**Figure 3.**
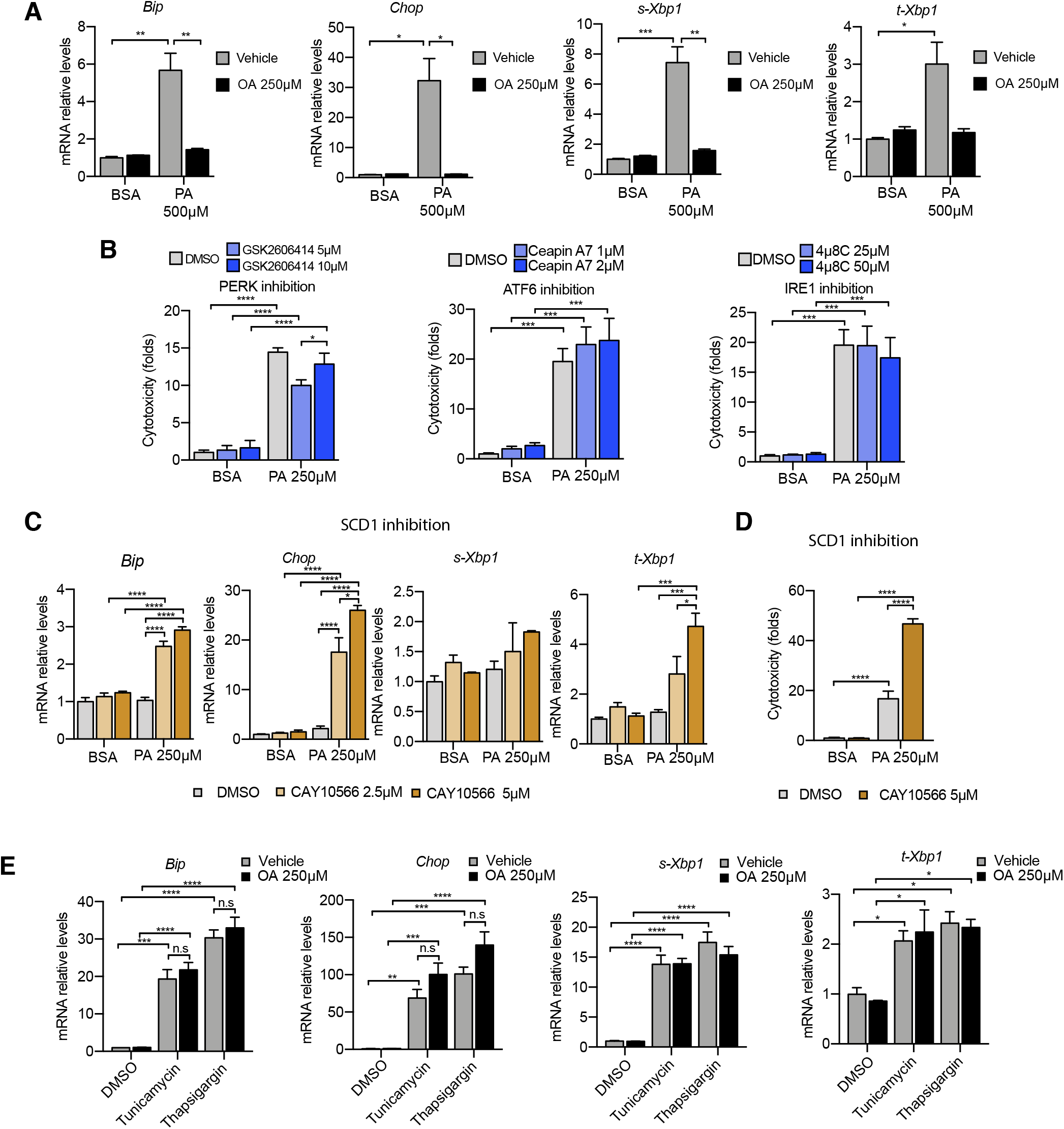
Excess of saturated fatty acids triggers the ER stress response. (A-C,E) Quantitative RT–PCR detection of ER stress markers in iRECs treated 16h with several combinations of BSA-fatty acids (A), PA plus ER stress inhibitors (B), PA plus SCD1 inhibitor (C) and Tunicamycin (10 µM) and Thapsigargin (1 µM) plus OA (D). (D) Cytotoxicity in iRECs at 24h treatment with PA 250µM plus the SCD1 inhibitor CAY10566. Fold representation of object counts/confluence. Data information: In (A-E) data are presented as mean ± SEM. *p<0.05, **p<0.01, ***p<0.001, ****p<0.0001, ns = non significant; two-way ANOVA and Holm-Sidak’s multiple comparisons test; n=3.

As *Scd1* and *Scd2* were top-regulated genes in the BSA-PA condition, we also measured ER stress markers and cytotoxicity in iRECs treated with a low dose of PA plus the chemical SCD inhibitor CAY10566. The inhibitor exposed the ER stress responses and enhanced the cytotoxicity induced by low-dose PA indicating that fatty acid desaturation is critical for preventing PA-induced ER stress (Fig 3C-D). We also measured ER stress markers in cells treated jointly with PA and etomoxir. Importantly, etomoxir exacerbated the ER stress response induced by PA, suggesting that decreased mitochondrial β-oxidation might increase the burden of PA in the ER (Fig S4A).

To investigate whether the protective effect of OA is specific for PA-induced ER stress, we treated BSA-OA cells with the chemical ER stressors tunicamycin and thapsigargin that are known to induce protein misfolding. Here, OA treatment did not show any protective effect against the ER stress caused by tunicamycin and thapsigargin. On the contrary, there was even a slight increase in the ER stress response, suggesting that OA might limit the capacity of the ER to deal with misfolded proteins (Fig 3E).

Finally, as prolonged ER stress can lead to the translocation of misfolded proteins into the cytoplasm via the ERAD system (Lemberg & Strisovsky, 2021), we reasoned that the increased expression of components of the autophagic machinery, such as p62, might be part of a proteostatic response. To confirm the presence of enhanced autophagy, we performed immunocytochemistry against the autophagy markers LC3 and p62. PA treatment induced the formation of LC3 puncta as well as big laminar structures positive for p62, all of which were cleared by the addition of OA (Fig S4B).

Altogether, the data demonstrate that ER stress (and autophagy as a consequence thereof) can be induced by PA *in vitro* and *in vivo*. Importantly, we also show that the protective effect of OA involves suppression of ER stress.

### PA increases membrane order in the ER

As previous studies have shown that ER stress can be induced by increased membrane order (Halbleib *et al*, 2017; Volmer *et al*, 2013), we next studied the effect of the fatty acid treatments on the ER membrane order. For this, we made use of C-Laurdan, an anisotropic dye that is able to visualize the degree of membrane order. After staining the cells, we segmented the perinuclear regions and quantified the generalized polarization (GP) values, which are indicative for membrane order. PA treatment decreased ER membrane fluidity (as evidenced by increased GP values) whereas OA increased it. The addition of OA to PA restored the GP values (Fig 4A-B) to normal levels. Thus, these results suggest that higher membrane order is associated with the PA-induced ER stress and that the mechanism by which OA suppresses ER stress involves restoring ER membrane order to homeostatic levels.

**Figure 4.**
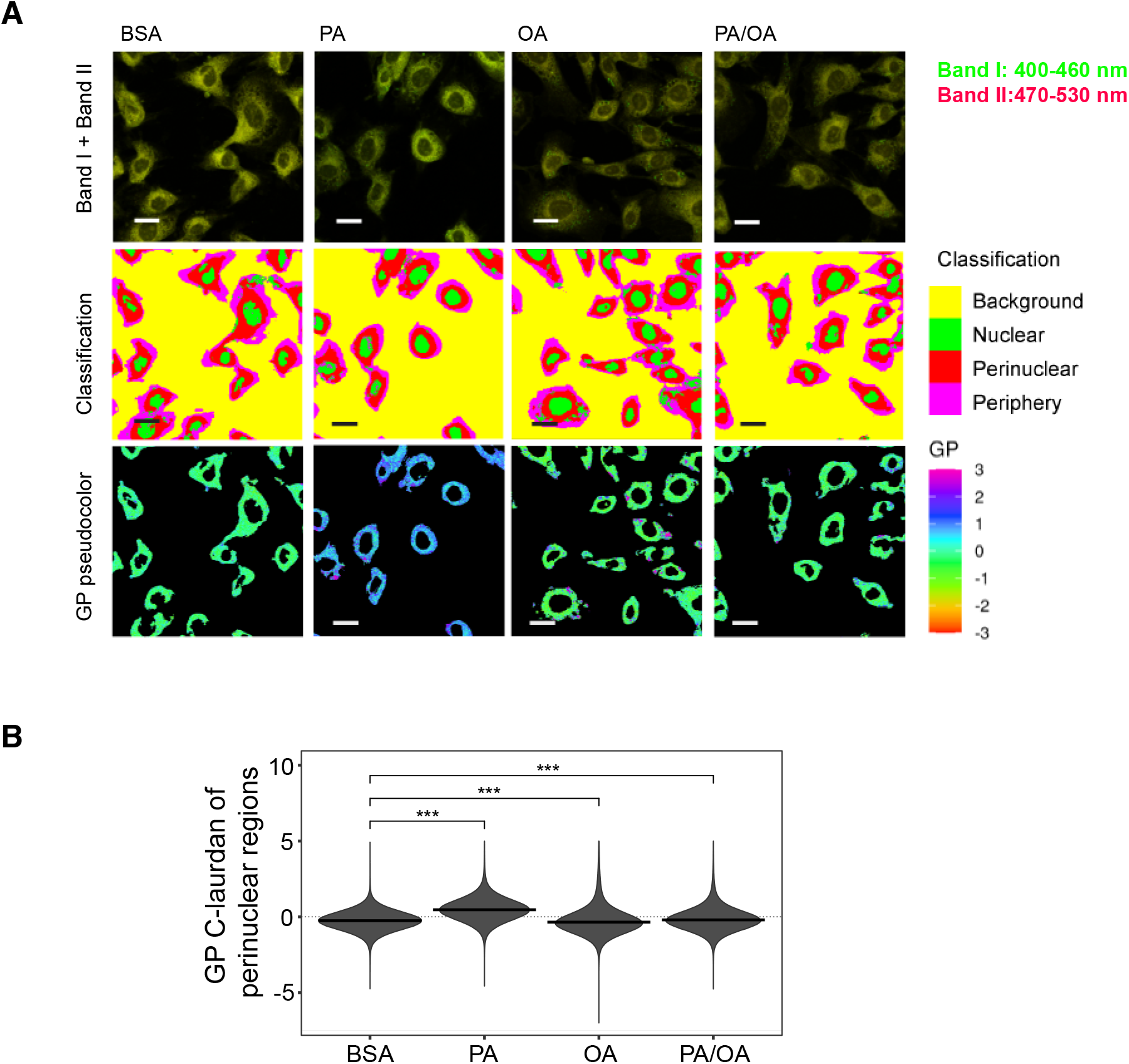
Perinuclear membrane order is increased by PA treatment and decreased by OA. (A) Representative C-Laurdan images of merged channels 1 and 2 (upper row), pixel classification (middle row) and GP pseudocoloured images. Membrane packing measurements by C-Laurdan in iRECs treated for 16h with BSA, PA 250µM, OA 250µM and PA 250µM plus OA 125µM. Scale bars: 20 μm. (B) GP values quantification of pixels classified as perinuclear from a single representative experiment. *p<0.05, **p<0.01, ***p<0.001, ****p<0.0001, ns = non significant ; Wilcoxon signed-rank test; n=3.

### PA impairs TAG synthesis and causes the accumulation of di-saturated TAG precursors and lysophospholipids

To identify the lipids that might mediate PA-induced changes in ER membrane order, iRECs treated with BSA, BSA-PA, BSA-OA and BSA-PA/OA were subjected to shotgun lipidomics and quantified. The results are represented schematically in Fig 5A-D. All four treatments did not differ much in the relative amount and degree of saturation of the major glycerophosholipid species (phosphatidylcholines, phosphatidylethanolamines, phosphatidylserines, phosphatidylglycerols and phosphatidylinositols). However, while OA treatment led to the strong formation of TAGs, PA-treated cells presented increased levels of precursors of TAGs synthesis, such as diacylglycerols, phosphatidic acid and lysophosphatidic acid. The addition of OA to PA enhanced the formation of TAGs, thereby reducing the accumulation of diacylglycerols, phosphatidic acid and lysophosphatidic acid. Moreover, PA treatment dramatically increased the saturation index of diacylglycerols, phosphatidic acid (Fig 5E-F) and lysophosphatidic acid, and, again, this was fully reverted by OA co-treatment. OA also decreased the concentration of free fatty acids, most likely as a consequence of the stimulated TAG synthesis (Fig 5G).

**Figure 5.**
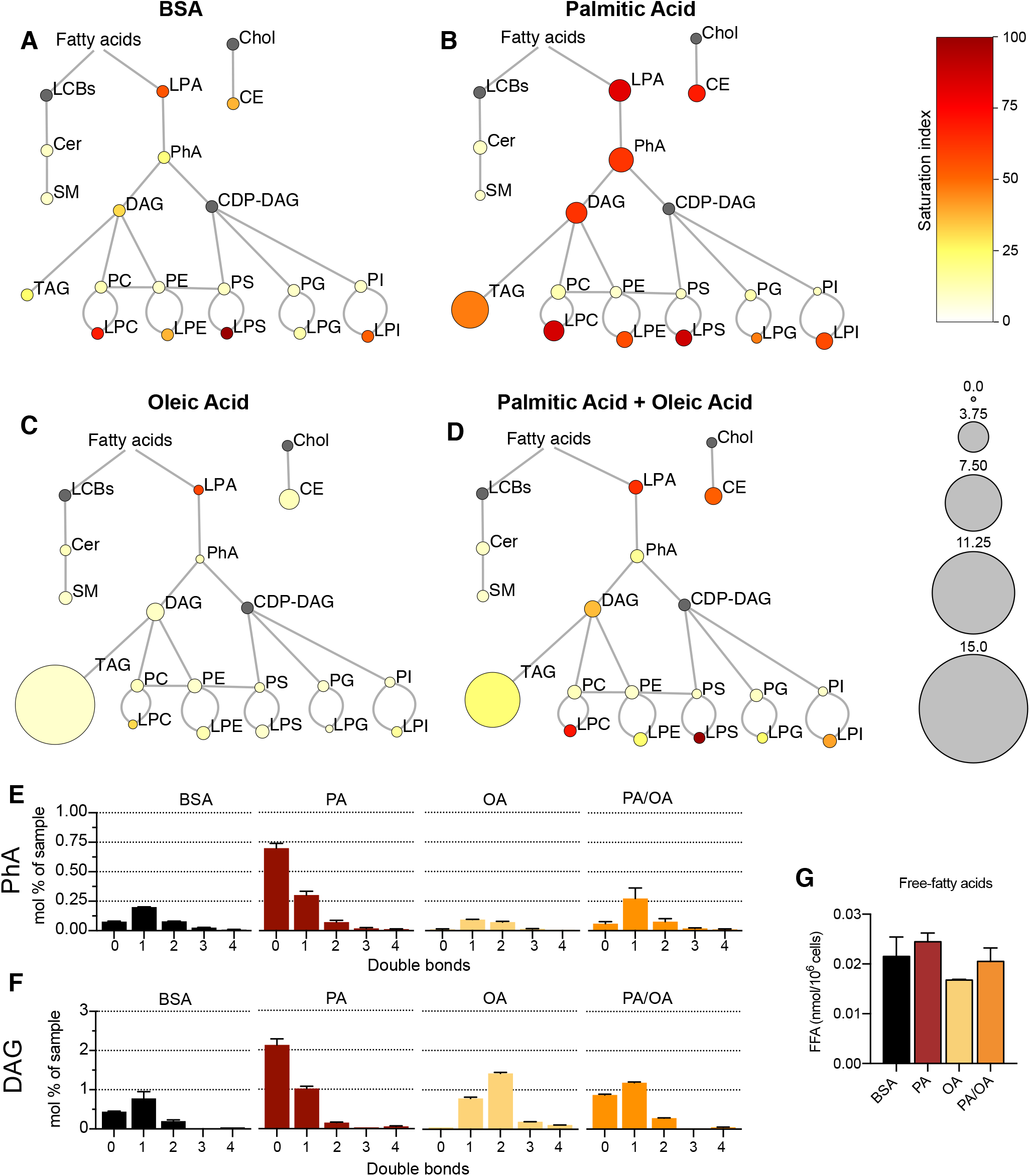
Lipidome of iRECs exposed to BSA-fatty acids. (A-D) Lipidome of iRECs treated for 16h with BSA (A), PA 250µM (B), OA 250µM (C) and PA 250µM plus OA 125µM (D). The scheme shows the relative levels of lipid classes presented as colour-coded circles. The lipid species were designated as saturated if all of its fatty acid chains were saturated, or unsaturated if it had at least one unsaturated fatty acid chain. The percentage of saturated lipid species is shown for each class from yellow (low saturation) to red (high saturation). Lipid classes not identified are shown in grey. Cholesterol is also presented in grey because it has no fatty acid chain. The size of the circles is set to the arbitrary unit of 1 for the BSA cells. G3P: glycerol-3-phosphate; LPA: lyso-phosphatidic acids; PhA: phosphatidic acids; DAG: diacylglycerol; TAG: triacylglycerol; PC: phosphatidylcholine: PE: phosphatidylethanolamine; LPE: lyso-phosphatidylethanolamine; LPC: lyso-phosphatidylcholine; PS: phosphatidylserine; LPS: lyso-phosphatidylserine; PI: phosphatidylinositol; LPI: lyso-phosphatidylinositol; PG: phosphatidylglycerol; LPG: lyso-phosphatidylglycerol; Cer: ceramide; SM: sphingomyelin; LCB: long-chain base; CDP: cytidine diphosphate; Chol: Cholesterol; CE: Cholesterol esthers. (n=3) (E-F) Relative amount of PhA (E) and DAG (F) species classified by the number of double bonds in iRECs treated for 16h with BSA, PA 250µM, OA 250µM and PA 250µM plus OA 125µM. Data are presented as mean ± SEM ; n=3. (G) Cytosolic free fatty acids in iRECs treated for 16h with BSA, PA 250µM, OA 250µM and PA 250µM plus OA 125µM. Data are presented as mean ± SEM ; One-way ANOVA and Holm-Sidak’s multiple comparisons test; n=3.

We also observed a general increase in lysophospholipid levels (lysophosphatidylcholines, lysophosphatidylethanolamines, lysophosphatidylserines, etc.) by PA that was accompanied by the transcriptional down- and upregulation *Lpcat1* and *Lpcat3*, respectively (Table S3). As part of the Lands’ cycle, LPCAT1 prefers palmitoyl-CoA (16:0-acyl-CoA) as an acyl donor to synthesize dipalmitoyl phosphatidylcholine, whereas LPCAT3 favors polyunsaturated FA-CoA as substrates (Wang & Tontonoz, 2019). The downregulation of LPCAT1 therefore likely reflects an attempt to dampen the production of oversaturated phosphatidylcholine species and, thereby, the lipid stress in the ER membrane. Interestingly, *Lpcat1* and *Lpcat3* are regulated in the reverse direction in OA-treated cells, demonstrating that a proper level of phosphatidylcholine unsaturation is critical for cell homeostasis (Table S3).

By computing biweight midcorrelation (bicor) (Song *et al*, 2012) between TF activities and lipids abundances, we also identified TF activities that associated with the TAG precursors. We revealed twenty-two TFs either negatively correlating with monounsaturated phosphatidic acid and diacylglycerol or positively correlating with di-saturated phosphatidic acids and diacylglycerols, or both (Fig S5A) as well as twelve TFs associating positively with di-saturated and negatively with monounsaturated lysophosphatidylcholine, lysophosphatidylethanolamine and lysophosphatidylinositol. Both lists include SREBF2 and HNF4A that have established roles in regulating many enzymes important for lipid metabolism (Fig S5B) (Guan *et al*, 2011; Madison, 2016; Wang & Tontonoz, 2019; Yin *et al*, 2011).

Altogether, our results support previous findings that, unlike MUFAs, SFAs impair TAG production in cultured cells (Listenberger *et al*, 2003; Piccolis *et al*, 2019). The accumulation of lysophosphatidic acid, phosphatidic acid and diacylglycerol further highlights the incapability of DGAT1 and DGAT2 to synthesize TAGs when too many acyl chains are saturated. Moreover, changes in the transcriptome reflect cellular attempts to maintain a proper level of phospholipid unsaturation, most likely to protect against ER stress. Finally, our multi-omic approach identified TFs potentially sensing ER membrane order and driving adaptation mechanisms.

### LD formation is stimulated by lipid unsaturation

As increased TAG synthesis allows for deposition of lipids in LDs, we next studied whether or not TAG levels correlate with LDs in iRECs. For this, we incubated the cells with BODIPY, a neutral lipid staining dye. OA treatment (0,25mM) resulted in both higher number and higher size of LDs compared to PA (0,25mM). Combined treatment (PA 0,25mM + OA 0,125mM) resulted in LDs smaller in size compared to OA alone (Fig 6A-C). Accordingly, the SCD1 inhibition blocked the formation of LDs in PA-treated cells resulting in LD numbers and sizes comparable to the BSA group (Fig 6A,D-E), suggesting that this enzyme (possibly together with SCD2) is required for the conversion of PA into MUFAs and subsequent LD formation. Moreover, performing a lipid trafficking study with the fatty acid analogue BODIPY-C12 showed that, when incubated only with BSA, the BODIPY-C12 dye was located in LDs but also in perinuclear membranes that co-localized with the ER marker calnexin. However, when incubated together with BSA-OA, BODIPY-C12 was strongly directed into LDs (Fig S6 A,B). In sum, MUFAs either imported from extracellular media or produced intracellularly by desaturases enhance the formation of TAGs and, subsequentially, ER-derived LDs.

**Figure 6.**
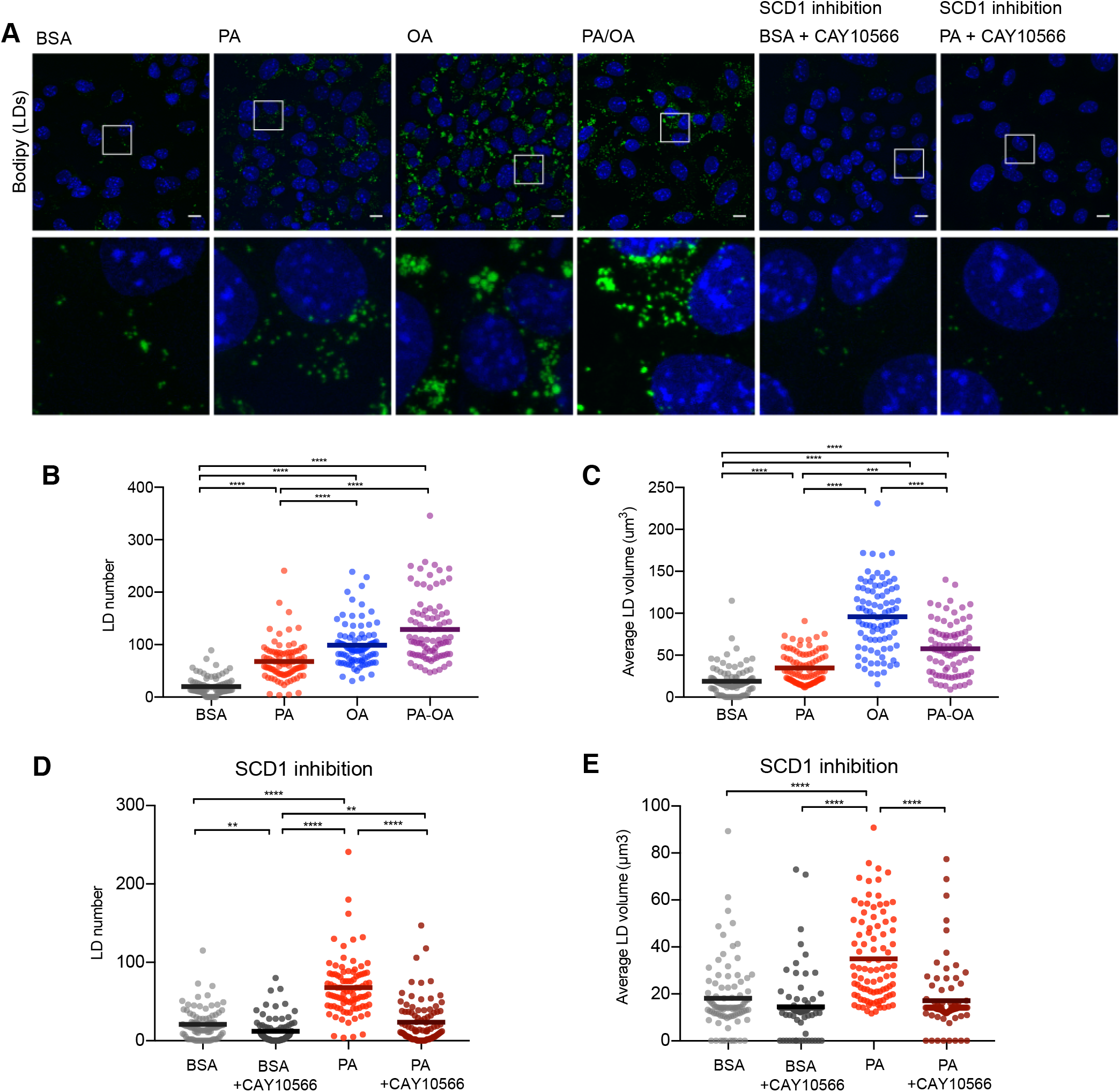
Saturated fatty acids impair the formation of lipid droplets. (A) Representative images of LDs stained using BODIPY in iRECs treated for 16h with BSA, PA 250µM, OA 250µM, PA 250µM plus OA 125µM, BSA plus the SCD1 inhibitor CAY10556(2.5µM) and PA plus CAY10556 (2.5µM). Scale bars: 10μm. (B-E) Quantification of LD number (B,D) and LD average volume (C,E) in iRECs treated for 16h with BSA, PA 250µM, OA 250µM, PA 250µM plus OA 125µM, BSA plus the SCD1 inhibitor CAY10556(2.5µM) and PA plus CAY10556 (2.5µM). Every dot represents the measurement in one single cell. Data information: In (B-E), data are presented as the mean + all values. *p<0.05, **p<0.01, ***p<0.001, ****p<0.0001 ; Kruskall-Wallis plus Dunn’s multiple comparisons test. (B-E) 10 cells per field from three fields were analysed for three independent biological replicates.

### Lipid droplets protect from PA-induced cytotoxicity

TAG formation is catalyzed by DGAT1 and DGAT2. To test their role in LD formation, we tested inhibitors of DGAT1 (T863) and DGAT2 (PF06424439) at different concentrations. We found that the combination of DGAT1 and DGAT2 inhibitors completely inhibited the formation of LDs (Fig S7A,B). Moreover, in iRECs treated with BSA or PA/OA, DGAT1/2 inhibitors caused only minimal changes in the relative amount and saturation of each lipid class (Fig 7A,C). By contrast, in iRECs treated with PA, the inhibitors caused major disturbances (Fig 7B). We observed a massive accumulation of oversaturated TAG precursors (lysophosphatidic acid, phosphatidic acid and diacylglycerols) as well as increased relative levels of saturated lysophospholipids. Especially remarkable is the increase of lysophosphatidylcholine, again reflecting the homeostatic response to prevent the production of oversaturated phosphatidylcholine species. A drastic decrease in cholesterol esters was also observed in all conditions treated with DGAT inhibitors, suggesting that TAG formation influences cholesterol esterification (Fig 7A-C).

**Figure 7.**
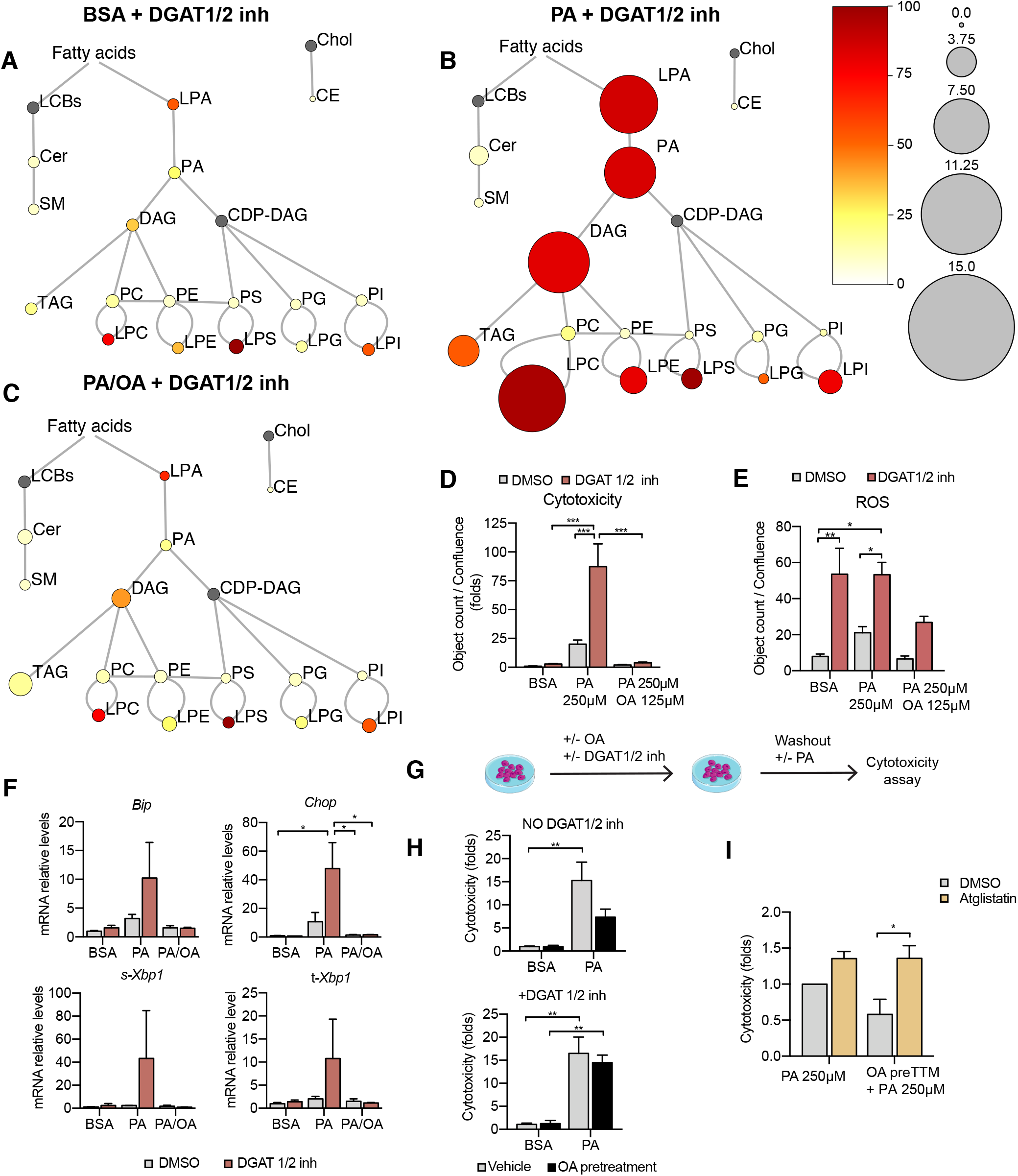
TAG synthesis protects from cytotoxic effects induced by exposure to saturated fatty acids. (A-C) Lipidome of iRECs treated for 16h with BSA + T863 30µM + PF 06424439 30µM, PA 250µM + T863 30µM + PF 06424439 30µM (B) and PA 250µM + OA 125µM + T863 30µM + PF 06424439 30µM (C). The scheme shows the relative levels of lipid classes presented as colour-coded circles. The lipid species were designated as saturated if all of its fatty acid chains were saturated, or unsaturated if it had at least one unsaturated fatty acid chain. The percentage of saturated lipid species is shown for each class from yellow (low saturation) to red (high saturation). Lipid classes not identified are shown in grey. Cholesterol is also presented in grey because it has no fatty acid chain. The size of the circles is set to the arbitrary unit of 1 for the BSA (Figure 4). G3P: glycerol-3-phosphate; LPA: lyso-phosphatidic acids; PhA: phosphatidic acids; DAG: diacylglycerol; TAG: triacylglycerol; PC: phosphatidylcholine: PE: phosphatidylethanolamine; LPE: lyso-phosphatidylethanolamine; LPC: lyso-phosphatidylcholine; PS: phosphatidylserine; LPS: lyso-phosphatidylserine; PI: phosphatidylinositol; LPI: lyso-phosphatidylinositol; PG: phosphatidylglycerol; LPG: lyso-phosphatidylglycerol; Cer: ceramide; SM: sphingomyelin; LCB: long-chain base; CDP: cytidine diphosphate; Chol: Cholesterol; CE: Cholesterol esthers. (n=3) (D) Cytotoxicity in iRECs at 36h treatment with BSA, PA 250µM and PA 250µM plus OA 125µM with or without the DGAT1/DGAT2 inhibitors T863/PF 06424439 (30µM). Fold representation of object counts/confluence. (E) Quantification of ROS generation in iRECs treated 16h with BSA, PA 250µM and PA 250µM plus OA 125µM with or without the DGAT1/DGAT2 inhibitors T863/PF06424439 (30µM). Data are presented as object count per well normalized by confluence. (F) Quantitative RT–PCR detection of ER stress markers in iRECs treated 16h with BSA, PA 250µM and PA 250µM plus OA 125µM with or without the DGAT1/DGAT2 inhibitors T863/PF06424439 (30µM). (G) Schematic representation of OA pretreatment plus PA insult experiment. (H) Cytotoxicity in iRECs at 36h after PA 250µM treatment. Cells were pre-treated for 16h with OA 500µM or BSA with or without DGAT1/DGAT2 inhibitors T863/PF06424439 (30µM). Fold representation of object counts/confluence. (I) Cytotoxicity in iRECs at 36h after PA 250µM treatment with or without atglistatin (25µM). Cells were pre-treated for 16h with OA 500µM or BSA. Fold representation of object counts/confluence. Data information: In (D,E,F,H,I) data are presented as mean ± SEM. *p<0.05, **p<0.01, ***p<0.001, ****p<0.0001; two-way ANOVA and Holm-Sidak’s multiple comparisons test. (A,B,C,D,F,H,I) n=3. (E) n=4.

Using the DGAT inhibitor combination, we could further show that PA-induced cytotoxicity was strongly exacerbated (Fig 7D). Accordingly, ROS and ER stress markers were increased upon DGAT inhibition (Fig 7E-F). Interestingly, the protective effect of OA concerning cytotoxicity and ER stress was unaltered in the presence of the DGAT inhibitors. With regard to ROS formation, the protective effect of OA was attenuated but did not reach statistical significance. Our results thus suggest that blocking LD formation through DGAT inhibition aggravates PA-induced cell death and ER stress because saturated TAG precursors accumulate in the ER membrane.

As LDs are dynamic organelles able to fine-tune the release of fatty acids in a lipolysis and re-esterification cycle (Chitraju *et al*, 2017), we hypothesized that MUFAs that are already stored into LDs could be released and help to channel PA towards TAG synthesis. To test this hypothesis, we pretreated the cells with OA 0.5mM with or without DGAT inhibitors and after washout, we challenged them with PA. Pretreatment with OA reduced PA-mediated cytotoxicity. When TAG synthesis was inhibited through DGAT inhibition, OA pretreatment did not show any effect on PA-induced cytotoxicity, suggesting that LDs rich in OA could protect from PA lipotoxicity (Fig 7G-H). To study the underlying mechanism, we used the ATGL inhibitor atglistatin to block the lipolytic release of fatty acids from LDs. Treatment with atglistatin in normal and starved conditions caused an increase in cell area occupied by LDs (Fig S7B,D). In order to test whether the release of fatty acids from LDs protects against PA-induced cytotoxicity, we pretreated cells with OA and exposed them to PA with and without the ATGL inhibitor. ATGL inhibition caused a significant increase in PA-induced cytotoxicity when compared to untreated cells (Fig 7I). Our results, therefore, suggest that LDs serve as a reservoir of MUFAs that can be released via lipolysis to buffer an overload of SFAs. MUFAs released from LDs could then facilitate the incorporation of SFAs into TAGs or decrease the packing density of ER membrane.

## Discussion

Lipids and lipid metabolites accumulate in tubules from humans and animal models of DKD, suggesting that lipotoxicity contributes to DKD pathogenesis (Herman-Edelstein *et al*., 2014; Kang *et al*., 2015). However, so far only limited data is available on the importance of dietary patterns for the progression of DKD in humans. While one study showed no protective effect of a Mediterranean diet compared to a low-fat control diet with regard to the DKD incidence in type 2 diabetics (Diaz-Lopez *et al*, 2015), two studies demonstrated renoprotective effects with adherence to Dietary Approaches to Stop Hypertension (DASH) and Mediterranean diets in cohorts of diabetic women (Jayedi *et al*, 2019; Yu *et al*, 2012).

In our mouse model, in which STZ-induced hyperglycemia was combined with a high fat diet, we found a depletion of the white adipose tissue that was accompanied by a significant weight gain in liver and kidney. Accordingly, the circulating lipids were increased and, in the kidney, lipid deposition was found in the tubular cells of the cortex. This suggests that, in addition to increased dietary fat intake, lipolysis in white adipose tissue might have contributed to renal fat accumulation. Strikingly, lipid deposition was primarily observed in the straight S2/S3 PTC segments while omitting glomeruli, S1 PTC segments and distal tubules. This lipid pattern could be explained by differences in fat uptake between the different nephron segments. As cubilin activity is known to be restricted to the S1 segment (Christensen *et al*, 2021; Ren *et al*, 2020), suggesting that other uptake pathways may account for lipid uptake in S2/S3. For example, it was recently shown that the scavenger receptor KIM1 mediates apical albumin-PA uptake in PTCs to promote DKD (Mori *et al*., 2021). What argues against this is that KIM1 was not upregulated by MUFA-HFD showing the strongest lipid accumulation. Another reason for the preferential lipid accumulation in S2/S3 could therefore be differences in lipid metabolism. Indeed, it is known that PTCs in the straight segments contain less mitochondria (Hall *et al*, 2009), possibly resulting in reduced consumption of lipid stores or LD buildup due to decreased β-oxidation. Also restricted expression of SCD1 to the straight S2/S3 segments could explain the lipid deposition pattern (Zhang *et al*, 2006). Upon cell entry, the intracellular fate and toxicity of the fatty acids clearly depended on the saturation of the acyl chain. While lipid storage in LDs was stronger in the MUFA-HFD treated mice than in SFA-HFD treated mice, tubular damage was reduced. This argues for a protective role of the MUFA diet in the tubules. However, as our mouse experiment was ended already 20 weeks after introducing the high fat diet, it would be interesting to study more long-term effects on renal function.

Using our cell culture model, we identified both mitochondrial and ER homeostasis as the main determinants of cell viability during PA-induced lipotoxic stress. The functional link between the two organelles was revealed by the finding that the inhibition of mitochondrial fatty acid uptake by etomoxir worsened ER stress and cytotoxicity induced by PA. When free SFAs cannot be oxidized in mitochondria they are incorporated into TAG precursors and lysophospholipids causing ER lipid bilayer stress. This could be suppressed by the addition of OA that promotes TAG and LD formation, which is also reflected by a reduction of free fatty acids by OA.

Due to the high degree of unsaturated lipids, the ER membrane is normally one of the most fluid membranes in the cell (Barelli & Antonny, 2016). Using Laurdan imaging, the ER membrane showed higher packing density upon PA treatment, most likely due to the accumulation of di-saturated TAG precursors, in particular diacylglycerols. This is in agreement with previous findings that di-saturated diacylglycerols are a poor substrate for DGAT activity and that the inhibition of GPAT enzymes that catalyze the first addition of fatty acids to the glycerol backbone is a promising approach for preventing lipotoxic cell injury (Piccolis *et al*., 2019). For diabetic kidneys, this might be particularly important as diacylglycerols have been implicated in insulin resistance due to their role in activating protein kinase C (PKC) isoforms (Lyu *et al*, 2020). How di-saturated diacylglycerols cause ER stress is not fully understood, but studies have shown that this likely involves PERK and IRE1-induced sensing of membrane lipid saturation (Halbleib *et al*., 2017; Volmer *et al*., 2013). Additionally, the altered ER membrane environment may lead to the misfolding of transmembrane proteins and subsequent induction of the unfolded protein response (UPR). Finally, the impairment of LD formation by PA may lower the capacity to buffer the lipotoxic stress in the ER.

Another PA effect we observed was the upregulation of the autophagic machinery possibly due to the appearance of protein aggregates caused by perturbations in the ER membrane. The activation of the autophagy machinery reflects the need to clear these misfolded protein aggregates. However, as the formation of the highly curved phagophore in the ER may be impaired by a high degree of phospholipid saturation (Kohler *et al*, 2009), it remains to be determined whether or not autophagy is successful at removing misfolded proteins or other cargoes. In particular, the accumulation of p62 suggests that this might not be the case. The induction of autophagy coupled with impaired execution might therefore be a vicious circle leading to additional ER stress.

The main finding of our study is that all observed PA-induced cytotoxic effects could be suppressed by adding OA. OA increased DGAT-mediated TAG which in turn facilitated LD formation. It is interesting that the inhibition of LD formation via the blocking of DGAT activity was dispensable for OA-mediated rescue effects in PA-induced stress. Only when cells were pre-treated with OA, then OA-mediated rescue effects were DGAT-dependent, suggesting that the preexistence of LDs is relevant for these rescue effects and/or that the beneficial effects of LD formation lag behind those associated with the formation of unsaturated phospholipids. As we also show that ATGL-mediated lipolysis is required for the OA rescue effect, our interpretation is that LDs can function as a reservoir for unsaturated lipids that can be released upon demand, for example when the ER membrane desaturation is increased. How this crosstalk between the ER and LDs could be regulated is an interesting question and should be subject of further studies.

In summary, we have undertaken a comprehensive analysis of PTC responses to lipotoxic stress. We find that the straight S2/S3 PTC segments are the primary site of fat deposition and that the damaging effects of renal fat deposition depend on the saturation of lipids. Our findings clearly point towards ER membrane saturation as a key determinant of cell viability that is regulated by LDs as a reservoir for MUFAs. Moreover, we identify transcriptional networks activated during PA-induced stress that can be used as a resource for a systems-level understanding of lipotoxic stress. As dietary effects on DKD progression are so far understudied in humans, our findings may provide new rationales for the prevention and management of DKD.

## Materials and Methods

### Animal experimentation and diets

All of the experimental protocols were performed with the approval of the animal experimentation ethics committee of the University Paris Descartes (CEEA 34), projects registered as 17-058 and 20-022. Mice were kept in a temperature-controlled room (22±1°C) on a 12/12 h light/dark cycle and were provided free access to commercial rodent chow and tap water prior to the experiments. The Monounsaturated Fatty Acid-High Fat Diet (MUFA-HFD) and Saturated Fatty Acid-High Fat Diet (SFA-HFD) were obtained from Research Diets. The HFDs (Research Diets, #D20072102 and #D20072103) had the following composition (in percentage of calories): 20% protein, 35% carbohydrates and 45% fat. The control diet (ENVIGO, #2018) composition was 24% protein, 58% carbohydrates and 18% fat. MUFA-HFD fat source was Olive Oil (95,1%) plus Soybean Oil (4,90%) and SFA-HFD fat source was Butter (95,1%) plus Soybean Oil (4,90%). HFDs were supplemented with Soybean oil to cover the essential need for polyunsaturated fatty acids (PUFAs). The detailed composition is shown in Table S1.

C57BL/6NCrl male 7-week-old mice were put on a control diet, MUFA-HFD or SFA-HFD with free access to food and water. At 11 weeks old, insulin deficiency was induced by intraperitoneal administration of streptozotocin (50 mg/kg per day for 5 consecutive days). Mice were fasted for 6h before streptozotocin (STZ) injections. STZ (Sigma-Aldrich, #S0130) was freshly prepared in 50 mM sodium citrate buffer pH 4.5 before administration. Control mice were injected with sodium citrate buffer. Blood samples were taken from the mandibular vein before STZ injections and every 4 weeks. Spot urine was collected at 10 and 14 weeks after STZ injection. Metabolic cages were avoided because they could have produced body weight loss and compromised the experiment. Food and water intake were measured per cage and values were divided by the number of mice in each cage.

The animals were sacrificed by cervical dislocation 16 weeks after STZ treatment. Blood was taken by intracardiac puncture and organs were perfused from the heart with PBS. The tissues (kidney, liver, heart, eWAT) were extracted, weighed and processed for histology and molecular analysis.

### Plasma and urine parameters

Urinary albumin and creatinine were determined by mouse albumin ELISA quantification kit (Bethyl Laboratories, #E99-134) and a home assay based on the Creatinine Parameter Assay Kit, (R&D Systems, #KGE005). Urine glucose levels were measured with a COBAS(r) 2000 analyzer (Roche). Blood glucose levels were measured using an “On Call GK dual” glucometer (Robe Medical, #GLU114). Blood levels of TAGs were measured by a colorimetric assay using the Triglyceride determination kit (Sigma-Aldrich, #MAK266). LCN2 plasma levels were determined by ELISA (R&D Systems, #AF1857-SP).

### Renal histopathology

Picro-Sirius Red, and Periodic Acid-Schiff (PAS) stainings were performed on paraffin-embedded sections. Oil Red O staining was performed on OCT frozen sections. Images were acquired in a slide scanner Nanozoomer HT2.0 C9600 (Hamatsu). Picro-Sirius Red images were analysed using Image J. Images were converted into RGB stack and the green channel was selected. The cortex region was segmented and the threshold was manually adjusted to determine the percentage of fibrotic area. Oil Red O staining was quantified using the pixel classification tool from QuPath.

### Cell culture

All cell lines were maintained at 37°C and 5% CO_2_. iRECS were cultured on 0.1% gelatin-coated flasks in Dulbecco’s modified Eagle’s medium (DMEM) (Lonza, #BE12-604F/U1) supplemented with penicillin/streptomycin (Sigma-Aldrich, #P4333), L-glutamine (Thermo Fisher Scientific, #25030024) and 10% (v/v) fetal bovine serum (FBS) (Thermo Fisher Scientific, #10270106). HK-2 cells were cultured in Renal Epithelial Cell Growth Medium (PromoCell, #C-26030) supplemented with SupplementMix (Promocell, #C-39606) for a final concentration of fetal calf serum 0,5% (v/v), FBS 1,5% (v/v), epidermal growth factor (10ng/mL), human recombinant insulin 5μg/mL, epinephrine 0,5 μg/mL, hydrocortisone 36 ng/mL, human recombinant transferrin 5g/mL, triiodo-L-thyronine 4 pg/mL. Primary mouse proximal tubule epithelial cells were isolated from mouse renal cortices as previously described (Legouis *et al*, 2015) and were cultured in Basal Medium 2 phenol red-free (PromoCell, #C-22216) supplemented in the same way as HK-2 cells. OK cells were cultured in DMEM/F12 (Thermo Fisher Scientific, #21331020) supplemented with penicillin/streptomycin, glutamine and 10% (v/v) fetal bovine serum (FBS).

### BSA-fatty acid conjugation

Fatty acid free-BSA (Sigma-Aldrich, #A8806) was added to complete medium to a final concentration of 1% (w/v), palmitic acid (Sigma-Aldrich, #P0500), oleic acid (Sigma-Aldrich #O1008) or a combination of both were added to the medium and incubated at 37°C for 30 minutes.

### Pharmacological treatments

TAG formation was inhibited using the DGAT1 inhibitor T863 (MedChemExpress, #HY-32219) and the DGAT2 inhibitor PF 06424439 (Bio-Techne, #6348/5). Import of fatty acids into mitochondria was blocked by CPT1 inhibitor etomoxir (Calbiochem, #236020). Desaturation of fatty acids was inhibited by the SCD1 inhibitor CAY10566 (MedChemExpress, #HY-15823-1mg). Lipolysis was inhibited by treating the cells with the ATGL inhibitor Atglistatin (Sigma-Aldrich, #SML1075). The ER stress response branches were inhibited individually using the PERK inhibitor GSK2606414 (MedChemExpress, #HY-18072), the IRE1a inhibitor 4μ8C (Sigma-Aldrich, #SML0949) and the ATF6 inhibitor Ceapin-A7 (Sigma-Aldrich, #SML2330). ER stress was chemically induced with tunicamycin (Sigma-Aldrich, #5045700001) and thapsigargin (Sigma-Aldrich, #586005). The concentration of each treatment is indicated in figures legends.

### RNAseq

Total RNA was isolated using the RNeasy Kit, (QIAGEN, #74104) including a DNAse treatment step. RNA quality was assessed by capillary electrophoresis using High Sensitivity RNA reagents with the Fragment Analyzer (Agilent Technologies) and the RNA concentration was measured by spectrophometry using the Xpose (Trinean) and Fragment Analyzer capillary electrophoresis.

RNAseq libraries were prepared starting from 1µg of total RNA using the Universal Plus mRNA-Seq kit (Nugen) as recommended by the manufacturer. The oriented cDNAs produced from the poly-A+ fraction were sequenced on a NovaSeq6000 from Illumina (Paired-End reads 100 bases + 100 bases). A total of ∼50 millions of passing filter paired-end reads was produced per library.

### Transcriptomics data processing and analysis

Galaxy platform was used to analyse the transcriptome data (Afgan *et al*, 2016). Quality check was assessed with FastQC v0.11.8 (http://www.bioinformatics.babraham.ac.uk/projects/fastqc/). After trimming with Trim Galore! v0.4.3 (http://www.bioinformatics.babraham.ac.uk/projects/trim_galore/), reads were aligned to the genome assembly GRCm38 using RNA STAR2 v2.5.2b (Dobin *et al*, 2013). Gene counts were calculated with featureCounts v1.6.4 (Liao *et al*, 2014) and differentially expressed gene (DEG) analysis was performed with DESeq2 v1.22.1 (Love *et al*, 2014). Gene counts were turned into log2 scale and used to perform Principal Components Analysis. Volcano plot for differences between PA and BSA treatments was drawn using EnchancedVolcano (https://github.com/kevinblighe/EnhancedVolcano). Significant DEGs were then divided into 12 clusters using the soft clustering tool Mfuzz v2.40.0 (Kumar & M, 2007) and computed clusters were assigned to expected expression change patterns. We investigated more closely the genes with expression patterns changed between BSA and PA treatment as well as PA and PA/OA treatment. Using Bioconductor package clusterProfiler v3.8.1 (Yu *et al*., 2012), enriched gene ontology categories for biological processes were compared and 25 most significant terms were plotted. All the analysis and data visualization was conducted in R v4.1.0.

For the transcription factor activity estimation, T-values of the differential analysis of PAvsCtrl, OAvsCtrl, OAvsPA/OA, PA/OAvsCtrl and PAvsPA/OA were used as input statistic for the viper algorithm (Alvarez *et al*, 2016). Regulon of TF and targets were obtained from the dorothea R package (Holland *et al*., 2020). Only TF-target interactions that belong in the confidence class A, B and C were kept. The viper function was used with default parameters, except minimum regulon size of 5, eset.filter parameter set to FALSE and pleiotropy parameter set to FALSE.

Biweight midcorrelation (bicor) was systematically estimated between TFs and lipid abundances using the bicor function of the WGCNA R package. To do so, average TF activities were first estimated at the level of individual conditions using the same method as described in the previous paragraph. Then, average TF activities across conditions (Ctrl, OA, PA/OA, PA) were correlated with average lipid abundance across the same conditions. To focus on the top associations between TFs and lipids, we selected TFs-lipids bicor coefficients above 0.97 or under -0.97. This coefficient value was chosen because it allowed us to select a number of associations that could be humanly investigated (e.i. less than 30). The full code for the analysis can be found in: https://github.com/saezlab/Albert_perez_RNA_lipid/tree/main/scripts

### Immunofluorescence and lipid droplet stainings

For immunocytochemistry, cells were washed three times in PBS, fixed for 20 min in 4% paraformaldehyde in PBS, blocked for 10 min in PBS + 3% BSA + 0,1% Tween + 0,1% Triton and incubated overnight at 4°C with primary antibodies diluted in PBS + 3% BSA + 0,1% tween + 0,1% triton. After washing, cells were incubated 2 h at room temperature with secondary antibodies (dilution 1:1000) and Hoechst (0.5 μg/mL) diluted in PBS + 3% BSA + 0,1% tween + 0,1% triton. Cells were washed three times in PBS and mounted in Roti®-Mount FluorCare (Roth, HP19.1). As primary antibodies guinea pig anti-p62 (1:1000, Progen, #GP62-C) and rabbit anti-LC3 (1:500, MBL International, #PM036) were used, and as secondary antibodies fluorescent conjugated Alexa Fluor 647 (Thermo Fisher Scientific, #A21244 and #A21450).

For lipid droplet imaging, BODIPY 493/503 (Thermo Fisher Scientific, #D3922) was incubated at 2,5 μg/mL together with secondary antibodies. For lipid trafficking studies, the fatty acid analog C1-BODIPY 500/510 C12 was incubated for 6h at a final concentration of 2μM (Thermo Fisher Scientific, #D3823). Images were acquired on a Leica TCS SP8 equipped with a 405-nm laser line and a white light laser with a 63x/1.4 DIC Lambda blue Plan Apochrome objective. Percentage of total cell area occupied by LDs was quantified with the ‘‘Analyze Particles’’-tool of Fiji on thresholded Z stack projections images of BODIPY 493/503. LD size and number were measured with Imaris by defining the volume of the LDs in 3D and then using the Spot Detector tool.

### Immunohistochemistry

For immunohistochemistry, kidney sections (4 μm) were generated from paraffin embedded tissue using a Leica RM2145 microtome and mounted on glass slides. Sections were first deparaffinized with xylene and rehydrated using a graded ethanol series. Antigen retrieval was performed by boiling the sections for 30min in TEG buffer (pH=9). Slides were washed in 50mM NM4Cl medium for 30min and then blocked with 1% BSA solution for 30min. Anti-KIM-1 antibody (RD Systems, #AF1817) was diluted (1:100) in 0.1% BSA, 0.3% Triton X-100. After overnight incubation at 4°C, slides were washed with 0.1% BSA and 0.3% Triton X-100 for three times. Next, they were incubated with Alexa Fluor 488 donkey anti-goat IgG (1:500; Invitrogen, #A-11055) for 1 hour at room temperature. Immunofluorescence images were taken by TCS SP5 confocal microscope at a resolution of 2048*2048 pixels.

### Membrane packing measurement by C-Laurdan

Membrane packing measurements by C-Laurdan imaging was performed as described previously (Kim *et al*, 2007; Levental *et al*, 2020). Briefly, cultured cells were fixed in 4% paraformaldehyde for 10 minutes. After two washes in PBS, cells were stained with 10 μg/mL C-Laurdan (kindly provided by Dr. B.R. Cho, Korea University, South Korea) in PBS for 15 minutes at room temperature. Subsequently, cells were imaged at 37°C by a confocal microscope equipped with a multiphoton laser (Zeiss LSM 780 NLO). C-Laurdan was excited by the 2-photon laser set to 800 nm. Two emission bands were acquired (band1: 43–463 nm, band2 473–503) using a 20x objective. Next, individual pixels (containing the signals of the two bands) were trained and classified using Ilastik (v. 1.3.3) (Berg *et al*, 2019) as ‘background’, ‘nuclear’, ‘perinuclear’, or ‘periphery’. Of all pixels except ‘background’, C-Laurdan Generalized Polarization (GP) values were calculated by the following formula: GP= (I_band1_-I_band2_)/(I_band1_ + I_band2_). For convenience, absolute GP values were *z*-scaled (mean = 0, sd = 1, over the full dataset). Images of pseudocolored GP values were generated in R (v. 4.1.0) using the packages ggplot2 (v. 3.3.5) and tiff (v. 0.1-8).

### Analysis of mRNA expression

Total RNA was isolated using the RNeasy Kit (74104, QIAGEN) including a DNAse treatment step. Concentration and purity of each sample was obtained from A260/A280 measurements in a micro-volume spectrophotometer NanoDrop-1000 (NanoDrop Technologies, Inc. Thermo Scientific). To measure the relative mRNA levels, quantitative (q)RT-PCR was performed using SYBR Green. cDNA was synthesized from 1 μg of total RNA with iScript™ cDNA Synthesis Kit (Bio-Rad, #1708891), following the manufacturer’s instructions. The Power SYBR® Green PCR Master Mix (Thermo Fisher Scientific, #4367659) was used for the PCR step. Amplification and detection were performed using the Mx3000P qPCR System (Agilent). Each mRNA from a single sample was measured in duplicate, using 18S and Beta-Actin as housekeeping genes. The primer sequences used were: 18S Fw 5’-CGGCTACCACATCCAAGGAA-3’, 18S Rv 5’-GCTGGAATTACCGCGGCT-3’, *β-Actin* Fw *5’-*GCTCTGGCTCCTAGCACCAT-3’, *β-Actin* Rv *5’-* GCCACCGATCCACACAGAGT-3’, *Bip* Fw *5’-*TTCAGCCAATTATCAGCAAACTCT-3’, *Bip* Rv *5’-* TTTTCTGATGTATCCTCTTCACCAGT-3’, *Chop* Fw *5’-*CCACCACACCTGAAAGCAGAA-3’, *Chop* Rv *5’-*AGGTGAAAGGCAGGGACTCA-3’, *Ccl5* Fw 5’-CCCTCACCATCATCCTCACT-3’, *Ccl5* Rv 5’-TCCTTCGAGTGACAAACACG-3’, *Fn1* Fw 5’-TTAAGCTCACATGCCAGTGC-3’, *Fn1* Rv 5’-TTAAGCTCACATGCCAGTGC-3’, *s-Xbp1 Fw 5’-*CTGAGTCCGAATCAGGTGCAG-3’, *s-Xbp1* Rv *5’-GTCCATGGGAAGATGTTCTGG-3’, t-Xbp1* Fw *5’-* TGGCCGGGTCTGCTGAGTCCG-3’, *t-Xbp1* Rv *5’-*GTCCATGGGAAGATGTTCTGG-3’.

### Cytotoxicity assay

An IncuCyte® S3 Live-Cell Analysis System was utilized to collect fluorescence and phase-contrast images over time. Cells were seeded at a density of 5000 cells per well in a 96 well culture microplate in the subconfluent experiments. For confluent experiments, 20000 cells per well were seeded. 24 hours post-plating cells, IncuCyte® Cytotox Red Reagent, (EssenBio, #4632) was added to the medium (dilution 1:2000) together with BSA-fatty acids. Cell confluence measurements and fluorescent objects were taken from 4 regions within each well and values were averaged to calculate mean confluence per well or object counts per well. Results were represented as object counts normalized by confluence.

### Mitochondrial membrane potential assay

Cells were seeded at a density of 5000 cells per well in a 96 well culture microplate. 8 hours post-plating cells, BSA-fatty acids treatment was performed. 24 hours post-plating cells, TMRE (tetramethylrhodamine, ethyl ester) (Thermo Fisher Scientific, # T669) was added to the medium (50nM) for 15 minutes. Cells were washed twice with PBS and full medium was added for the fluorescence measurement. Cells were immediately imaged in the red channel of IncuCyte® S3 Live-Cell Analysis System. Results were represented as the image’s average of the objects’ mean fluorescent intensity.

### ROS detection assay

Cells were seeded at a density of 5000 cells per well in a 96 well culture microplate. 24 hours post-plating cells, DHE (Dihydroethidium) (Thermo Fisher Scientific, #D23107) was added to the medium (10μM) together with BSA-fatty acids. 4 hours after DHE addition, cells were imaged in the red channel of IncuCyte® S3 Live-Cell Analysis System. Cell confluence measurements and fluorescent objects were taken from 4 regions within each well and values were averaged to calculate mean confluence per well or object counts per well. Results were represented as objects counts normalized by confluence.

### Lipidomics

#### Lipid extraction for mass spectrometry lipidomics

Mass spectrometry-based lipid analysis was performed by Lipotype GmbH (Dresden, Germany) as described (Sampaio *et al*, 2011). Lipids were extracted using a two-step chloroform/methanol procedure (Ejsing *et al*, 2009). Samples were spiked with internal lipid standard mixture containing: cardiolipin 14:0/14:0/14:0/14:0 (CL), ceramide 18:1;2/17:0 (Cer), diacylglycerol 17:0/17:0 (DAG), hexosylceramide 18:1;2/12:0 (HexCer), lyso-phosphatidate 17:0 (LPA), lyso-phosphatidylcholine 12:0 (LPC), lyso-phosphatidylethanolamine 17:1 (LPE), lyso-phosphatidylglycerol 17:1 (LPG), lyso-phosphatidylinositol 17:1 (LPI), lyso-phosphatidylserine 17:1 (LPS), phosphatidate 17:0/17:0 (PhA), phosphatidylcholine 17:0/17:0 (PC), phosphatidylethanolamine 17:0/17:0 (PE), phosphatidylglycerol 17:0/17:0 (PG), phosphatidylinositol 16:0/16:0 (PI), phosphatidylserine 17:0/17:0 (PS), cholesterol ester 20:0 (CE), sphingomyelin 18:1;2/12:0;0 (SM), sulfatide d18:1;2/12:0;0 (Sulf), triacylglycerol 17:0/17:0/17:0 (TAG) and cholesterol D6 (Chol). After extraction, the organic phase was transferred to an infusion plate and dried in a speed vacuum concentrator. 1st step dry extract was re-suspended in 7.5 mM ammonium acetate in chloroform/methanol/propanol (1:2:4, V:V:V) and 2nd step dry extract in 33% ethanol solution of methylamine in chloroform/methanol (0.003:5:1; V:V:V). All liquid handling steps were performed using Hamilton Robotics STARlet robotic platform with the anti-droplet control feature for organic solvents pipetting.

### MS data acquisition

Samples were analyzed by direct infusion on a QExactive mass spectrometer (Thermo Scientific) equipped with a TriVersa NanoMate ion source (Advion Biosciences). Samples were analyzed in both positive and negative ion modes with a resolution of Rm/z=200=280000 for MS and Rm/z=200=17500 for MSMS experiments, in a single acquisition. MSMS was triggered by an inclusion list encompassing corresponding MS mass ranges scanned in 1 Da increments (Surma *et al*, 2015). Both MS and MSMS data were combined to monitor CE, DAG and TAG ions as ammonium adducts; PC, PC O-, as acetate adducts; and CL, PA, PE, PE O-, PG, PI and PS as deprotonated anions. MS only was used to monitor LPA, LPE, LPE O-, LPI and LPS as deprotonated anions; Cer, HexCer, SM, LPC and LPC O-as acetate adducts and cholesterol as ammonium adduct of an acetylated derivative (Liebisch *et al*, 2006).

### Data analysis and post-processing

Data were analysed with in-house developed lipid identification software based on LipidXplorer (Herzog *et al*, 2012; Herzog *et al*, 2011). Data post-processing and normalization were performed using an in-house developed data management system. Only lipid identifications with a signal-to-noise ratio >5, and a signal intensity 5-fold higher than in corresponding blank samples were considered for further data analysis.

Lipidomics data were represented using the cytoscape software (https://cytoscape.org).

### Intracellular FFA measurements

iRECs were seeded at a density of 1.5 x10^6^ in a 10cm plate. Free fatty acids were measured by a colorimetric assay using the Free fatty acid quantification Kit (Sigma-Aldrich, #MAK044) following manufacturer’s instructions.

### Statistics

Statistical analyses were performed using R Studio and GraphPad Prism. P value less than 0.05 was considered significant (*p < 0.05, **p < 0.01, ***p < 0.001, ****p < 0.0001). More details on statistics can be found in each figure legend.

## Acknowledgements

We thank Sophie Berissi for help with histology staining procedures, Hermann-Josef Gröne for help with the histological evaluation, Shrey Kohli and Berend Iserman for help with urine analysis and Yung-Hsin Shin and Gwenn Le Meur for excellent technical assistance. We acknowledge the European Research Council (ERC) under the European Horizon 2020 research and innovation programme (Grant agreement No. 865408 (RENOPROTECT) to M.S and No. 804474 (DiRECT) to S.S.L), the Deutsche Forschungsgemeinschaft (DFG SI1303/5-1 (Heisenberg-Programm) and DFG SI1303/6-1), the Steno Collaborative Grant from the NovoNordisk Foundation (NNF18OC0052457) (all to M.S.) and the Swiss National Science Foundation (NCCR Kidney.CH and 310030_189102) to S.S.L. A.P.-M. was funded by a postdoctoral fellowship from the Fondation pour la Recherche Médicale (FRM-SPF20170938629).

## Author contributions

**Albert Pérez-Martí:** conceptualization, formal analysis, investigation, writing - original draft, visualization, project administration.

**Suresh Ramakrishnan:** investigation.

**Jiayi Li:** investigation.

**Aurelien Dugourd:** software, formal analysis, data curation and visualization.

**Martijn R. Molenaar:** investigation, formal analysis and visualization

**Luigi R. De La Motte:** investigation.

**Kelli Grand:** software, formal analysis and visualization.

**Anis Mansouri:** software, formal analysis and visualization.

**Mélanie Parisot:** investigation.

**Soeren S. Lienkamp:** resources and funding acquisition.

**Julio Saez-Rodriguez:** resources and funding acquisition.

**Matias Simons:** conceptualization, writing - original draft, project administration, supervision, funding acquisition.

## Data and materials availability

All data are available in the main text or the supplementary materials.

