## Supplemental Information for "Increasing triacylglycerol formation and lipid storage by unsaturated lipids protects renal proximal tubules in diabetes"

Fig S1

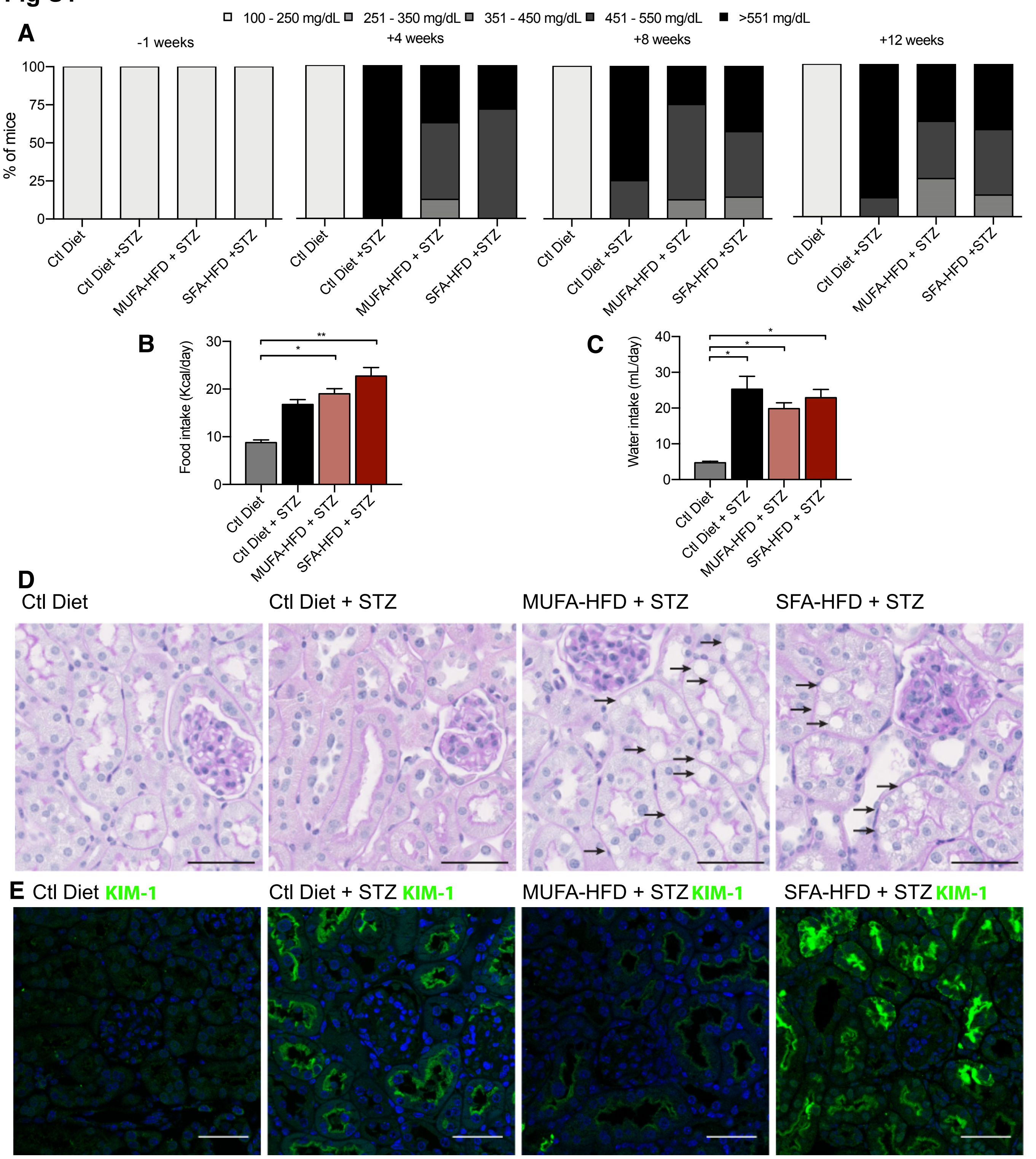

A
100 - 250 mg/dL
251 - 350 mg/dL
351 - 450 mg/dL
451 - 550 mg/dL
>551 mg/dL
-1 weeks
+4 weeks
+8 weeks
+12 weeks
% of mice
100
75
50
25
0
Ctl Diet
Ctl Diet +STZ
MUFA-HFD + STZ
SFA-HFD +STZ
B
Food intake (Kcal/day)
30
20
10
0
*
**
Ctl Diet
Ctl Diet + STZ
MUFA-HFD + STZ
SFA-HFD + STZ
C
Water intake (mL/day)
40
30
20
10
0
*
*
*
D
Ctl Diet
Ctl Diet + STZ
MUFA-HFD + STZ
SFA-HFD + STZ
E
Ctl Diet KIM-1
Ctl Diet + STZ KIM-1
MUFA-HFD + STZ KIM-1
SFA-HFD + STZ KIM-1

**Fig S2**

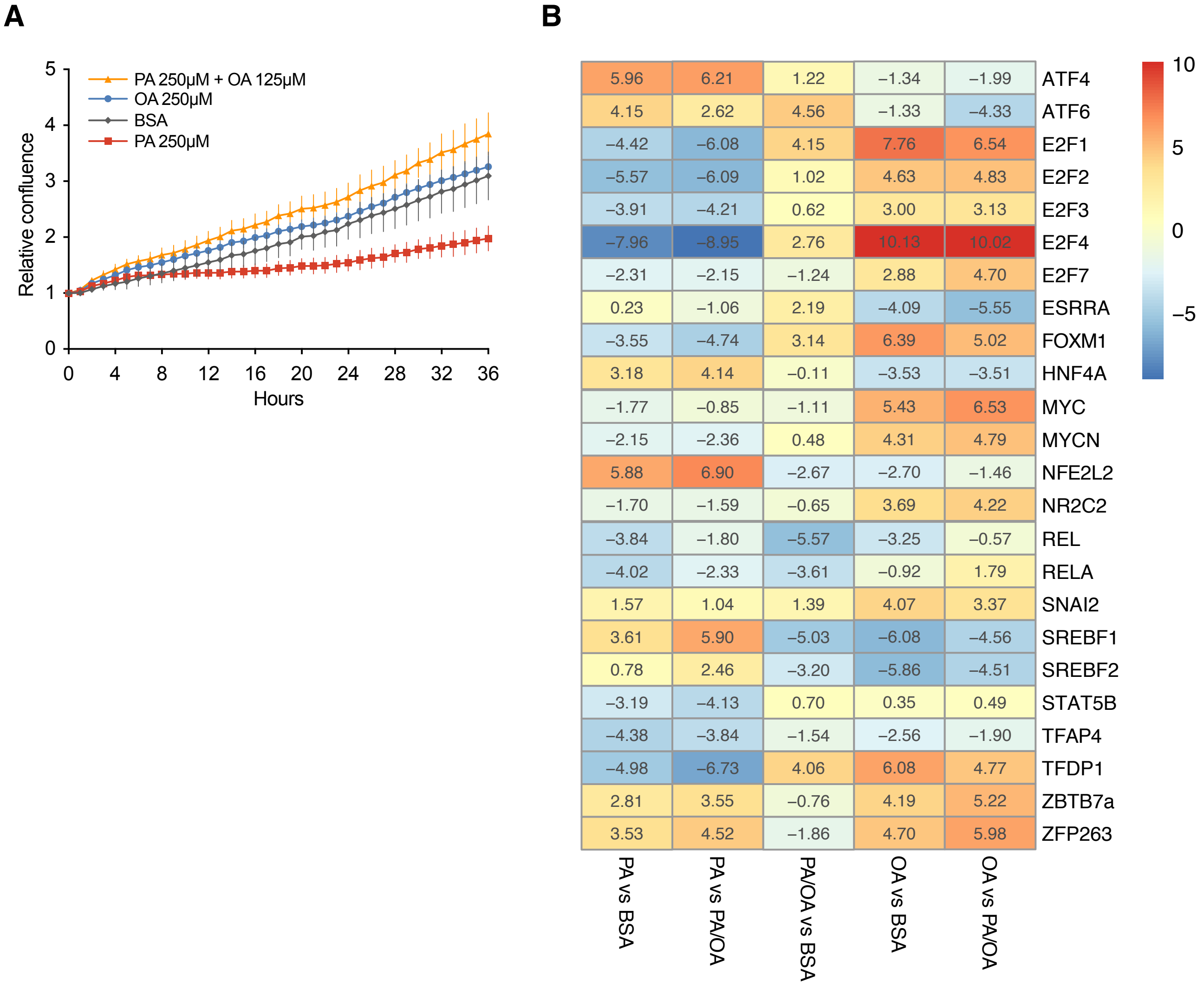
A
B
PA 250μM + OA 125μM
OA 250μM
BSA
PA 250μM
Relative confluence
Hours
0 1 2 3 4 5
0 4 8 12 16 20 24 28 32 36
ATF4 5.96 6.21 1.22 −1.34 −1.99
ATF6 4.15 2.62 4.56 −1.33 −4.33
E2F1 −4.42 −6.08 4.15 7.76 6.54
E2F2 −5.57 −6.09 1.02 4.63 4.83
E2F3 −3.91 −4.21 0.62 3.00 3.13
E2F4 −7.96 −8.95 2.76 10.13 10.02
E2F7 −2.31 −2.15 −1.24 2.88 4.70
ESRRA 0.23 −1.06 2.19 −4.09 −5.55
FOXM1 −3.55 −4.74 3.14 6.39 5.02
HNF4A 3.18 4.14 −0.11 −3.53 −3.51
MYC −1.77 −0.85 −1.11 5.43 6.53
MYCN −2.15 −2.36 0.48 4.31 4.79
NFE2L2 5.88 6.90 −2.67 −2.70 −1.46
NR2C2 −1.70 −1.59 −0.65 3.69 4.22
REL −3.84 −1.80 −5.57 −3.25 −0.57
RELA −4.02 −2.33 −3.61 −0.92 1.79
SNAI2 1.57 1.04 1.39 4.07 3.37
SREBF1 3.61 5.90 −5.03 −6.08 −4.56
SREBF2 0.78 2.46 −3.20 −5.86 −4.51
STAT5B −3.19 −4.13 0.70 0.35 0.49
TFAP4 −4.38 −3.84 −1.54 −2.56 −1.90
TFDP1 −4.98 −6.73 4.06 6.08 4.77
ZBTB7a 2.81 3.55 −0.76 4.19 5.22
ZFP263 3.53 4.52 −1.86 4.70 5.98
PA vs BSA
PA vs PA/OA
PA/OA vs BSA
OA vs BSA
OA vs PA/OA
10
5
0
−5

Fig S3

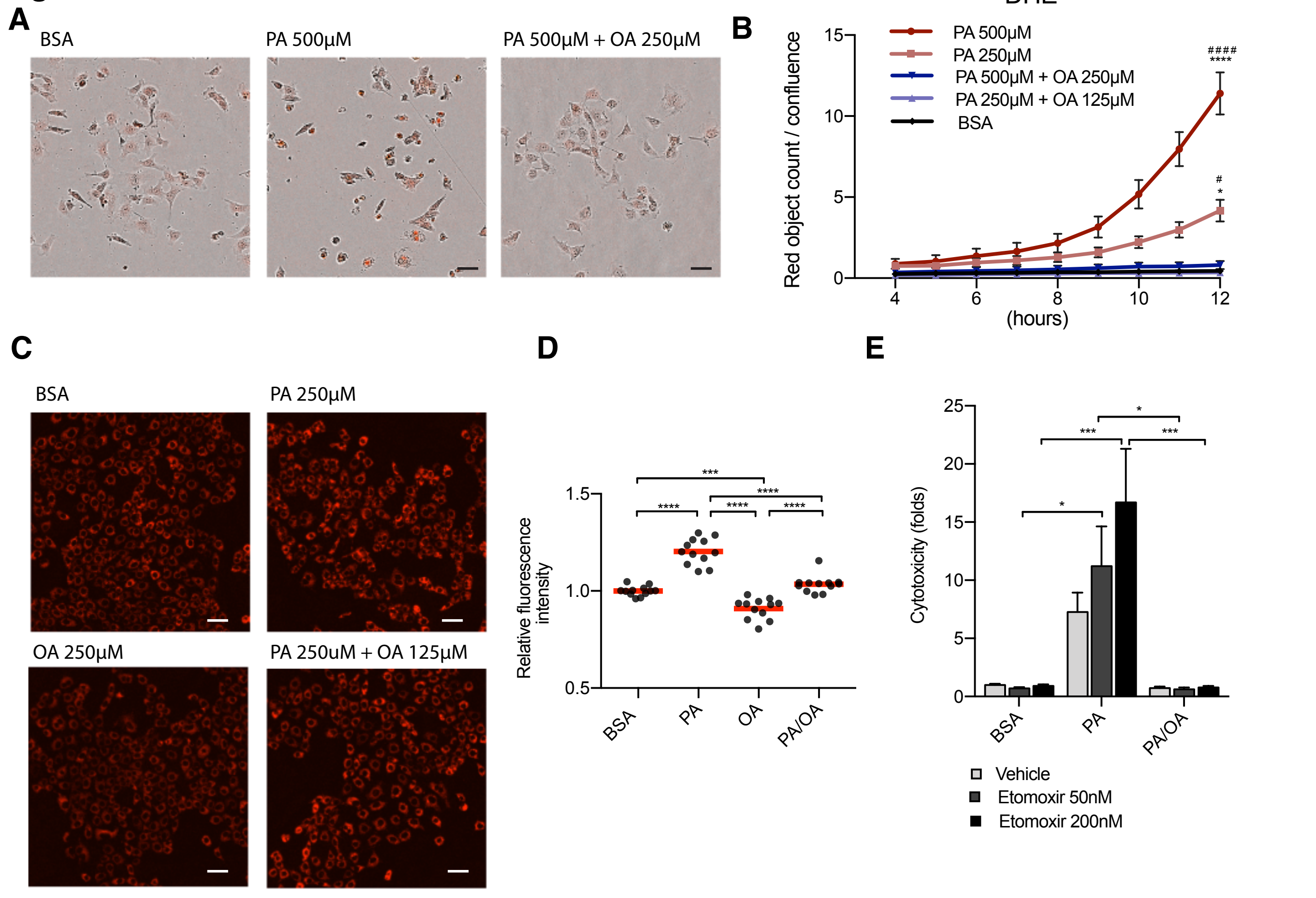

A
BSA
PA 500μM
PA 500μM + OA 250μM
B
DHE
Red object count / confluence
PA 500μM
PA 250μM
PA 500μM + OA 250μM
PA 250μM + OA 125μM
BSA
####
****
#
*
15
10
5
0
4
6
8
10
12
(hours)
C
BSA
PA 250μM
OA 250μM
PA 250uM + OA 125μM
D
Relative fluorescence intensity
1.5
1.0
0.5
***
****
****
****
****
BSA
PA
OA
PA/OA
E
Cytotoxicity (folds)
25
20
15
10
5
0
*
***
***
*
BSA
PA
PA/OA
Vehicle
Etomoxir 50nM
Etomoxir 200nM

**Fig S4**

**A**

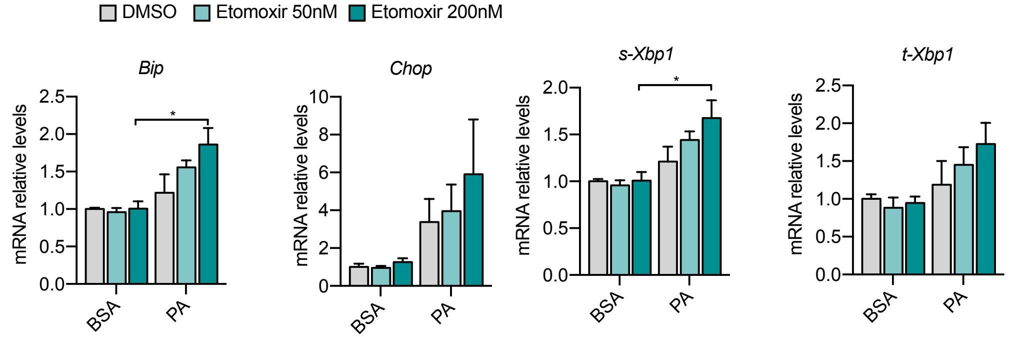

DMSO
Etomoxir 50nM
Etomoxir 200nM
Bip
Chop
s-Xbp1
t-Xbp1
mRNA relative levels
BSA
PA
*

**B**

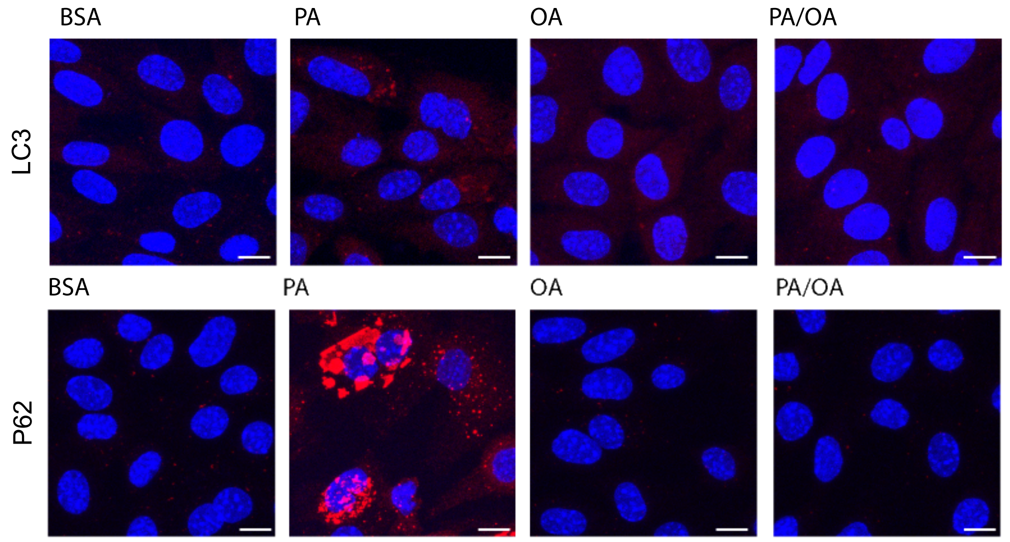

BSA
PA
OA
PA/OA
LC3
P62

### Fig S5

**A** **B**

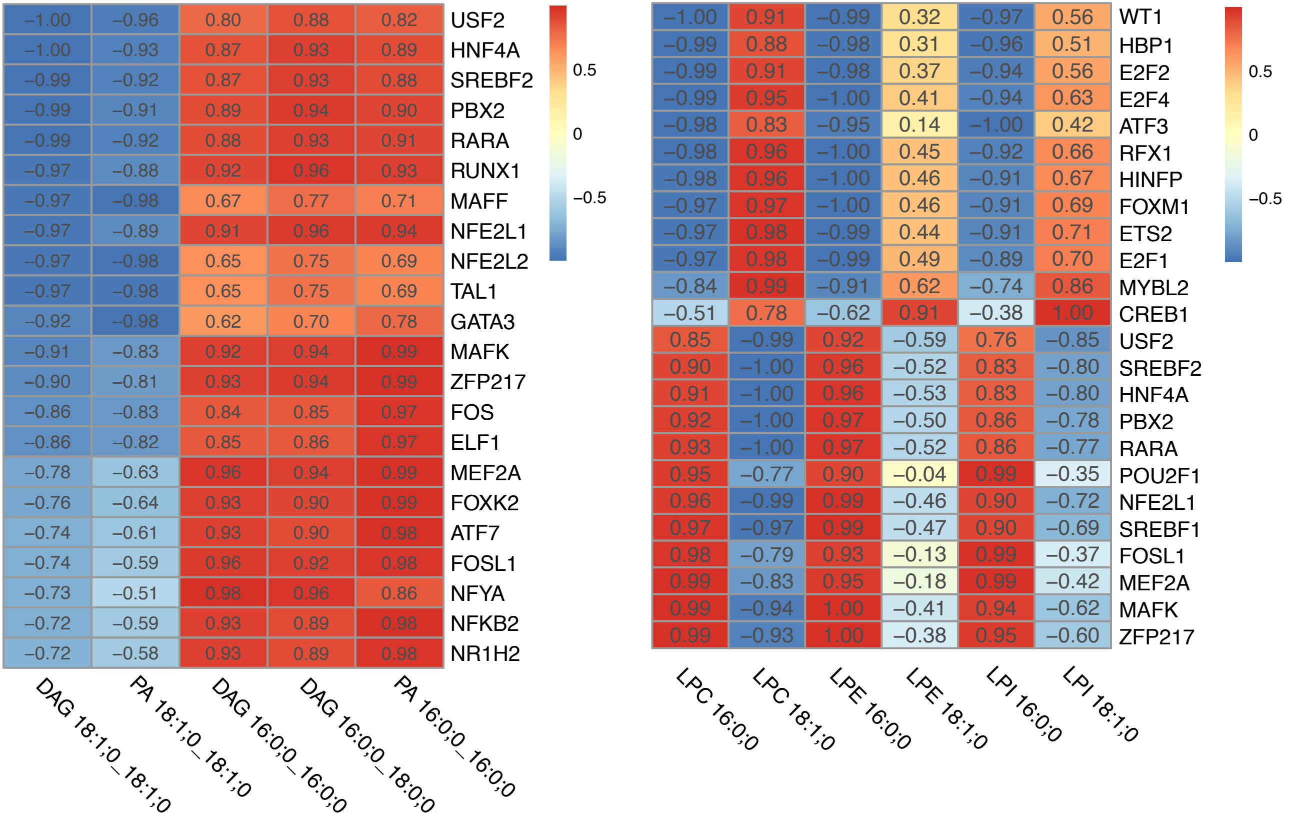
A
−1.00 −0.96 0.80 0.88 0.82 USF2
−1.00 −0.93 0.87 0.93 0.89 HNF4A
−0.99 −0.92 0.87 0.93 0.88 SREBF2
−0.99 −0.91 0.89 0.94 0.90 PBX2
−0.99 −0.92 0.88 0.93 0.91 RARA
−0.97 −0.88 0.92 0.96 0.93 RUNX1
−0.97 −0.98 0.67 0.77 0.71 MAFF
−0.97 −0.89 0.91 0.96 0.94 NFE2L1
−0.97 −0.98 0.65 0.75 0.69 NFE2L2
−0.97 −0.98 0.65 0.75 0.69 TAL1
−0.92 −0.98 0.62 0.70 0.78 GATA3
−0.91 −0.83 0.92 0.94 0.99 MAFK
−0.90 −0.81 0.93 0.94 0.99 ZFP217
−0.86 −0.83 0.84 0.85 0.97 FOS
−0.86 −0.82 0.85 0.86 0.97 ELF1
−0.78 −0.63 0.96 0.94 0.99 MEF2A
−0.76 −0.64 0.93 0.90 0.99 FOXK2
−0.74 −0.61 0.93 0.90 0.98 ATF7
−0.74 −0.59 0.96 0.92 0.98 FOSL1
−0.73 −0.51 0.98 0.96 0.86 NFYA
−0.72 −0.59 0.93 0.89 0.98 NFKB2
−0.72 −0.58 0.93 0.89 0.98 NR1H2
DAG 18:1;0_18:1;0
PA 18:1;0_18:1;0
DAG 16:0;0_16:0;0
DAG 16:0;0_18:0;0
PA 16:0;0_16:0;0
0.5
0
−0.5
B
−1.00 0.91 −0.99 0.32 −0.97 0.56 WT1
−0.99 0.88 −0.98 0.31 −0.96 0.51 HBP1
−0.99 0.91 −0.98 0.37 −0.94 0.56 E2F2
−0.99 0.95 −1.00 0.41 −0.94 0.63 E2F4
−0.98 0.83 −0.95 0.14 −1.00 0.42 ATF3
−0.98 0.96 −1.00 0.45 −0.92 0.66 RFX1
−0.98 0.96 −1.00 0.46 −0.91 0.67 HINFP
−0.97 0.97 −1.00 0.46 −0.91 0.69 FOXM1
−0.97 0.98 −0.99 0.44 −0.91 0.71 ETS2
−0.97 0.98 −0.99 0.49 −0.89 0.70 E2F1
−0.84 0.99 −0.91 0.62 −0.74 0.86 MYBL2
−0.51 0.78 −0.62 0.91 −0.38 1.00 CREB1
0.85 −0.99 0.92 −0.59 0.76 −0.85 USF2
0.90 −1.00 0.96 −0.52 0.83 −0.80 SREBF2
0.91 −1.00 0.96 −0.53 0.83 −0.80 HNF4A
0.92 −1.00 0.97 −0.50 0.86 −0.78 PBX2
0.93 −1.00 0.97 −0.52 0.86 −0.77 RARA
0.95 −0.77 0.90 −0.04 0.99 −0.35 POU2F1
0.96 −0.99 0.99 −0.46 0.90 −0.72 NFE2L1
0.97 −0.97 0.99 −0.47 0.90 −0.69 SREBF1
0.98 −0.79 0.93 −0.13 0.99 −0.37 FOSL1
0.99 −0.83 0.95 −0.18 0.99 −0.42 MEF2A
0.99 −0.94 1.00 −0.41 0.94 −0.62 MAFK
0.99 −0.93 1.00 −0.38 0.95 −0.60 ZFP217
LPC 16:0;0
LPC 18:1;0
LPE 16:0;0
LPE 18:1;0
LPI 16:0;0
LPI 18:1;0
0.5
0
−0.5

### Fig S6

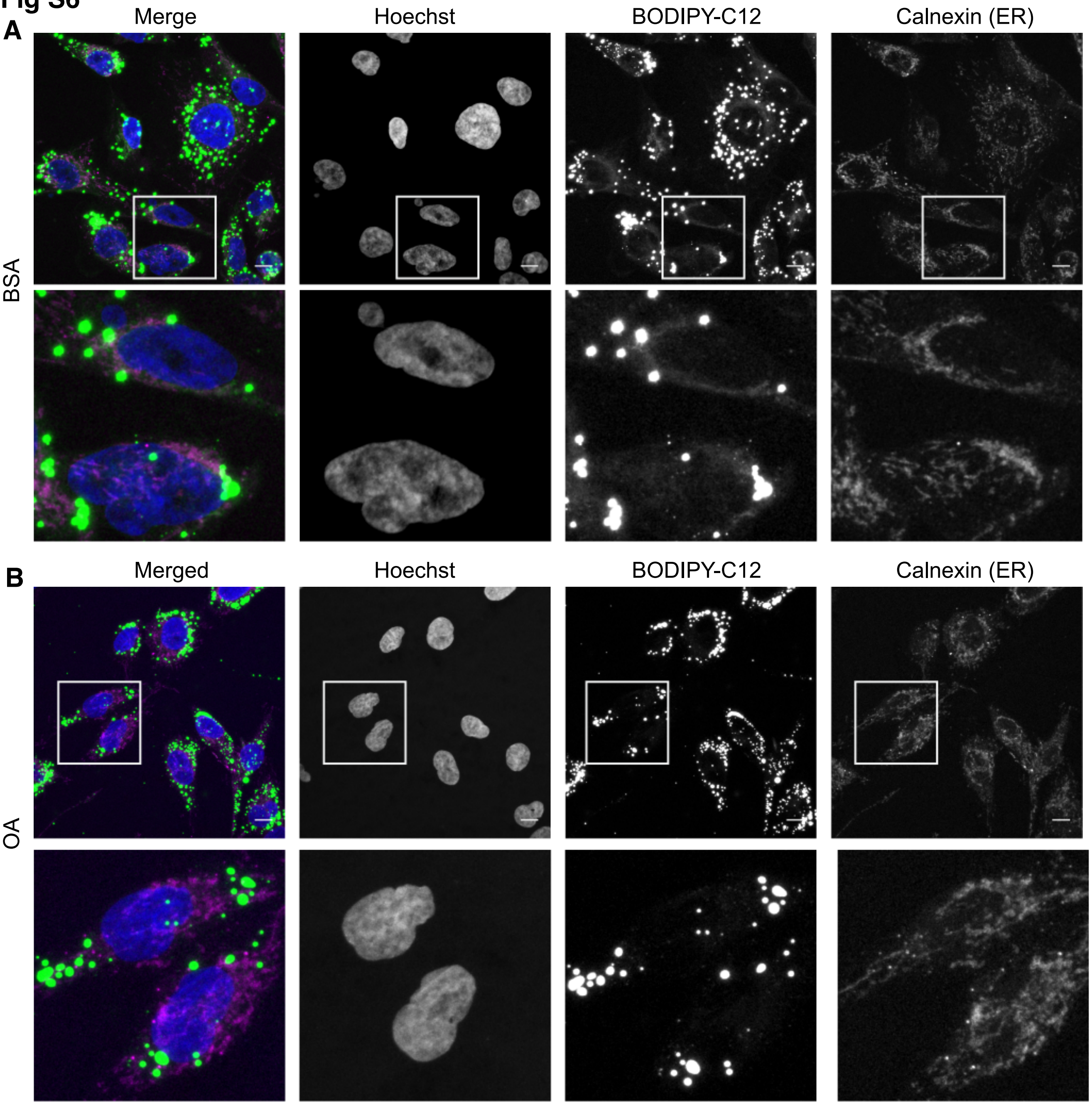

A
Merge
Hoechst
BODIPY-C12
Calnexin (ER)
BSA
B
Merged
Hoechst
BODIPY-C12
Calnexin (ER)
OA

Fig S7

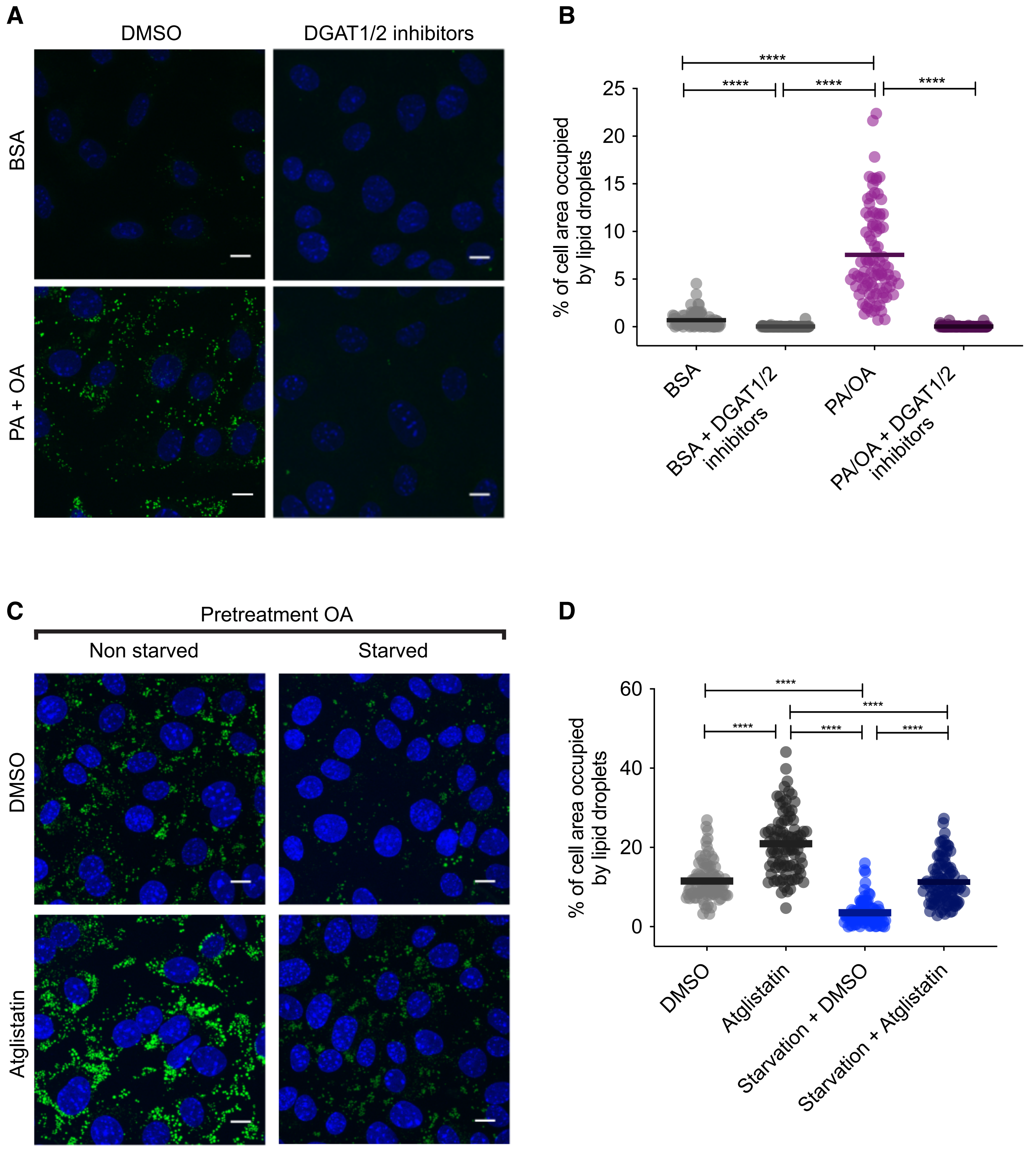
A
DMSO
DGAT1/2 inhibitors
BSA
PA + OA
B
****
****
****
****
% of cell area occupied by lipid droplets
25
20
15
10
5
0
BSA
BSA + DGAT1/2 inhibitors
PA/OA
PA/OA + DGAT1/2 inhibitors
C
Pretreatment OA
Non starved
Starved
DMSO
Atglistatin
D
****
****
****
****
****
% of cell area occupied by lipid droplets
60
40
20
0
DMSO
Atglistatin
Starvation + DMSO
Starvation + Atglistatin

**Supplemental Figure Legends**

**Figure S1 Diabetic phenotypes in STZ-injected mice**

(A) Non-fasted blood glucose levels 1 week before and 4, 8 and 12 weeks after STZ injection.

(B) Daily food intake 9 weeks after STZ injection.

(C) Daily water intake 9 weeks after STZ injection.

(D) Representative bright-field images of mouse kidney cortex PAS staining. Arrows indicate lipid droplets. Scale bars: 50*μm.*

(E) Representative confocal fluorescence images of mouse kidney cortex stained for KIM-1. Scale bars: 50*μm.*

Data information: In (B,C) data are presented as mean ± SEM. *p<0.05, **p<0.01; one-way ANOVA plus Holm-Sidak's multiple comparisons test. In (A) n=7 Ctl diet, n=8 Ctl Diet + STZ, n=8 MUFA-HFD + STZ, n=7 SFA-HFD + STZ. In (B,C) measurements were performed on mice grouped by cages; n=2 Ctl diet, n=2 Ctl Diet + STZ, n=2 MUFA-HFD + STZ, n=4 SFA-HFD + STZ.

**Figure S2 Cell growth and estimated TF activities in response to PA, OA and PA/OA**

(A) Relative change in confluence of iRECs treated for 36h with BSA, PA 250μM, OA 250μM and PA 250μM + OA 125μM; data are presented as mean ± SEM; n=3.

(B) TFs activity estimated from the transcriptome of iRECs treated for 16h with BSA, PA 250μM, OA 250μM and PA 250μM + OA 125μM; n=3.

**Figure S3 OA suppresses PA-induced oxidative stress**

(A) Representative images of ROS generation in iRECs visualised by the incorporation of DHE into the nucleus at 12h. Images are the combination of bright-field and fluorescence. Scale bars: 50*μm.*

(B) Quantification of ROS generation in iRECs treated with several BSA-fatty acids combinations. Data are presented as object count per well normalized by confluence.

(D,E) Representative fluorescence images (D) of iRECs incubated with TMRE after 16h treatment with BSA, PA 250µM, OA 250µM and PA 250µM + OA 125µM and its quantification (E). Scale bars: 50*µm.*

Data information: In (B,C), data are presented as mean ± SEM. In (E), data are presented as mean and the value from every replicate is presented as a dot. *p<0.05, **p<0.01, ***p<0.001, ****p<0.0001. (B,D) one-way ANOVA and Holm-Sidak's multiple comparisons test, (E) two-way ANOVA and Holm-Sidak's multiple comparisons test. (B,C) n=3. (E) 4 technical replicates from three independent experiments were pooled

**Figure S4 Mitochondrial fatty acid uptake and autophagy during PA-induced ER stress**

(A) Quantitative RT–PCR detection of ER stress markers in iRECs treated 16h with BSA and PA 250µM with or without the etomoxir. *p<0.05; two-way ANOVA and Holm-Sidak's multiple comparisons test; n=3.

(B) Representative images of LC3 and P62 immunostainings of iRECs treated for 16h with BSA, PA 250µM and PA 250µM plus OA 125µM; n=3. Scale bars: 10µm.

**Figure S5 Correlation of TF activity with relevant components of the ER membrane**

(A) TFs with higher correlation score respect relevant saturated (PA 16:0_16:0, DAG 16:0_16:0, DAG 16:0_18:0) and unsaturated (PA 18:1_18:1, DAG18:1_18:1) TAG precursors. TFs with FPKM<1 were omitted.

(B) TFs with higher correlation score respect relevant saturated (LPC 16:0, LPE 16:0, LPI 16:0) and unsaturated (LPC 18:1, LPE 18:1, LPI 18:1) lysophospholipids. TFs with FPKM<1 were omitted.

**Figure S6 OA channels a fatty acid analog into lipid droplets**

Representative images of HK-2 cells treated for 6h with BODIPY-C12 together with BSA (A) or BSA-OA 250µM (B) and co-stained with the ER marker calnexin; n=3 for BODIPY C-12 staining and n=1 for co-staining with calnexin. Scale bars: 10µm.

**Figure S7 Pharmacological inhibition of lipid droplets biogenesis and degradation**

(C,D) Representative images (C) and quantification (D) of LDs stained using BODIPY in iRECs cultured for 16h in starvation medium (1% FBS) and complet medium (10% FBS + OA 500 µM) with or without atglistatin (25µM). Cells were pre-treated for 16h with OA 500µM.

Data information: In (B,D), data are presented as the mean + all values. *p<0.05, **p<0.01, ***p<0.001, ****p<0.0001 ; Kruskall-Wallis plus Dunn's multiple comparisons test. (B,D) 10 cells per field from three fields were analysed for three independent biological replicates. Every dot represents the measurement in one single cell.

**Supplemental Tables**

**Table S1: Nutrient composition of mouse diets**

| | CTL DIET | MUFA-HFD | SFA-HFD |
|---|---|---|---|
| Macronutrients | | | |
| Carbohydrates (kcal%) | 58 | 35 | 35 |
| Protein (kcal%) | 24 | 20 | 20 |
| Fat (kcal%) | 18 | 45 | 45 |
| Energy density (kcal/g) | 3.1 | 4.52 | 4.52 |
| Source of fat | | | |
| Soybean Oil (kcal%) | 18 | 2.2 | 2.2 |
| Olive Oil (kcal%) | 0 | 42.8 | 0 |
| Butter, Anhydrous (kcal%) | 0 | 0 | 42.8 |
| Saturated | 15.5 | 14.3 | 62.6 |
| Monounsaturated | 23.9 | 69.5 | 30.7 |
| Polyunsaturated | 60.6 | 15.9 | 6.8 |
| Typical fatty acids composition (%) | Soybean Oil | Olive Oil | Butter, Anhydrous |
| C4, Butyric | 0 | 0 | 3.2 |
| C6, Caproic | 0 | 0 | 1.9 |
| C8 Caprylic | 0 | 0 | 1.1 |
| C10, Capric | 0 | 0 | 2.5 |
| C12, Lauric | 0 | 0 | 2.8 |
| C14, Myristic | 0.1 | 0 | 10 |
| C14:1, Myristoleic | 0 | 0 | 1.5 |
| C16, Palmitic | 10.4 | 11.5 | 26.2 |
| C16:1, Palmitoleic | 0.1 | 1.2 | 2.3 |
| C18, Stearic | 3.9 | 2.3 | 12.1 |
| C18:1, Oleic | 23 | 70.5 | 25.1 |
| C18:2, Linoleic | 51.8 | 13.0 | 2.3 |
| C18:3, Linolenic | 7.4 | 0.6 | 0 |
| C20, Arachidic | 0.4 | 0.4 | 1 |
| C20:1 | 0 | 0.2 | 0 |
| C22, Behenic | 0.3 | 0 | 0 |
| C24, Lignoceric | 0.2 | 0 | 0 |

**Table S2: Urinary parameters**

| | Ctl Diet | Ctl Diet + STZ | MUFA-HFD + STZ | SFA-HFD + STZ |
|---|---|---|---|---|
| UACR (µg/mg) | 28.71 ± 8.30 | 64.81 ± 38.28 | 177 ± 59.10 **** # | 97.41 ± 21.76* |
| Glucose / Creatinine (mg/mg) | 0.8599 ± 0.34 | 2734 ± 1360*** | 1433 ± 859.90* | 1435 ± 821.10* |

Urine albumin-to-creatinine ratio (UACR) and Urine glucose-to-creatinine ratio at 14 weeks after STZ injection. Data are presented as mean ± StDev. *p<0.05, ****p<0.0001 vs Ctl Diet; #p<0.05 vs Ctl Diet + STZ (Kruskal-Wallis plus Dunn's multiple comparisons test).
